# Post-translational control of PlsB is sufficient to coordinate membrane synthesis with growth in *Escherichia coli*

**DOI:** 10.1101/728451

**Authors:** Marek J Noga, Ferhat Büke, Niels JF van den Broek, Nicole Imholz, Nicole Scherer, Flora Yang, Gregory Bokinsky

**Author notes:** Translational Metabolic Laboratory, Department of Laboratory Medicine, Radboudumc, Nijmegen, The Netherlands (Marek J Noga).

## Abstract

Every cell must produce enough membrane to contain itself. However, the mechanisms by which the rate of membrane synthesis is coupled with the rate of cell growth remain unresolved. By comparing substrate and enzyme concentrations of the fatty acid and phospholipid synthesis pathways of *Escherichia coli* across a 3-fold range of carbon-limited growth rates, we show that the rate of membrane phospholipid synthesis during steady-state growth is determined principally through allosteric control of a single enzyme, PlsB. Due to feedback regulation of the fatty acid pathway, PlsB activity also indirectly controls synthesis of lipopolysaccharide, a major component of the outer membrane synthesized from a fatty acid synthesis intermediate. Surprisingly, concentrations of the enzyme that catalyses the committed step of lipopolysaccharide synthesis (LpxC) do not vary across steady-state growth conditions, suggesting that steady-state lipopolysaccharide synthesis is modulated primarily via indirect control by PlsB. In contrast to steady-state regulation, we find that responses to environmental perturbations are triggered directly via changes in acetyl-CoA concentrations, which enables rapid adaptation. Adaptations are further modulated by ppGpp, which regulates PlsB activity during slow growth and growth arrest. The strong reliance of the membrane synthesis pathway upon post-translational regulation ensures both reliability and responsiveness of membrane synthesis.

**Significance:** How do bacteria cells grow without breaking their membranes? Although the biochemistry of fatty acid and membrane synthesis is well-known, how membrane synthesis is balanced with growth and metabolism has remained unclear. This is partly due to the many control points that have been discovered within the membrane synthesis pathways. By precisely establishing the contributions of individual pathway enzymes, our results simplify the model of membrane biogenesis in the model bacteria species *Escherichia coli*. Specifically, we find that allosteric control of a single enzyme, PlsB, is sufficient to balance growth with membrane synthesis and to ensure that growing *E. coli* produces sufficient membrane. Identifying the signals that activate and deactivate PlsB will answer the question of how membrane synthesis is synchronized with growth.

## Introduction

All cells build and expand their membranes at a pace that must be coordinated with growth, as excess or insufficient membrane production can be fatal. In Gram-negative bacteria, construction of the double membrane from fatty acid precursors demands coordination between phospholipid (PL) and lipopolysaccharide (LPS) synthesis pathways (1, 2) as well as with protein synthesis, which supplies the lipoproteins that tether the outer membrane to the peptidoglycan cell wall (3). Many elements of membrane synthesis regulation that act on either transcription or enzymes activities have been proposed or identified. However, how each of these individual elements are used to regulate membrane synthesis during steady-state growth remains unclear (4).

The question of how membrane construction is coordinated with growth can be approached by considering how the metabolic pathways that synthesize membrane building blocks (PL and LPS) are regulated. Biosynthetic fluxes are regulated either by control of enzyme concentrations or by direct control of enzyme activity. Examples of both forms of regulation can be readily found: in *Escherichia coli*, the steady-state protein synthesis rate is controlled by ribosome concentration, which is transcriptionally regulated to balance amino acid supply with translation demand (5). In contrast, fluxes through central carbon metabolism are controlled instead by concentrations of substrates and inhibitors (i.e. post-translational control) rather than by changes in enzyme concentration (6, 7). Post-translational control is also known to contribute to membrane synthesis regulation (8–13), however, how cells use transcriptional and post-translational control to coordinate membrane synthesis with growth has never been well-defined.

In order to understand how the model Gram-negative species *Escherichia coli* coordinates membrane synthesis with growth, we quantified substrates and enzymes of the fatty acid and PL synthesis pathways in both steady-state and dynamic conditions. We find that post-translational control of the first enzyme in the PL synthesis pathway, PlsB, is sufficient to ensure steady-state regulation of PL synthesis. In contrast, transcriptional regulation maintains stable enzyme concentrations regardless of growth rate. Furthermore, due to feedback regulation of the fatty acid pathway, PlsB exerts strong control over LPS synthesis via its effects on concentrations of an LPS fatty acid precursor.

## Results

### The PL to biomass ratio varies inversely with μ

Lipid precursors of both LPS and PL are synthesized in the cytosol as fatty acyl thioesters covalently attached to acyl carrier protein (acyl-ACP). Fatty acid synthesis is initiated by carboxylation of acetyl-CoA by the acetyl-CoA carboxylase complex (ACC) to produce malonyl-CoA, from which the malonyl group is transferred to holo-ACP by FabD. Malonyl-ACP is used to elongate fatty acids iterated cycles beginning with condensation with acyl-ACP, reduction, dehydration, and reduction. The first step in PL synthesis is catalysed by the membrane-bound enzyme PlsB, which synthesizes lysophosphatidic acid (LPA) from long-chain acyl-ACP and *sn*-glycerol-3 phosphate (G3P). G3P is produced either from glyceraldehyde-3-phosphate, an intermediate of central carbon metabolism, or via glycerol catabolism **(Figure 1A)**. We grew cultures of *E. coli* NCM3722 in 6 defined media that support a 3-fold range of growth rates (μ). Quantities of major PL species phosphatidylglycerol (PG), phosphatidylethanolamine (PE), and cardiolipin (CL) per unit biomass (as determined by culture optical density (OD) (14)) decrease slightly with increasing μ **(Figure 1B)**, consistent with previous observations in *E. coli* and other bacteria (15, 16). The PG/PE ratio remains constant across all μ. We quantified steady-state PL flux by multiplying total PE LC/MS counts by μ, which closely approximates the total PL synthesis rate as PE turnover is slow relative to synthesis (17) and PG/PE ratio is stable. PL flux increases by 2-fold as μ increases by 3-fold **(Figure 1C)**. The higher PL to biomass ratio of slow-growing cells likely reflects their higher surface-to-volume ratio (18).

**Figure 1.**
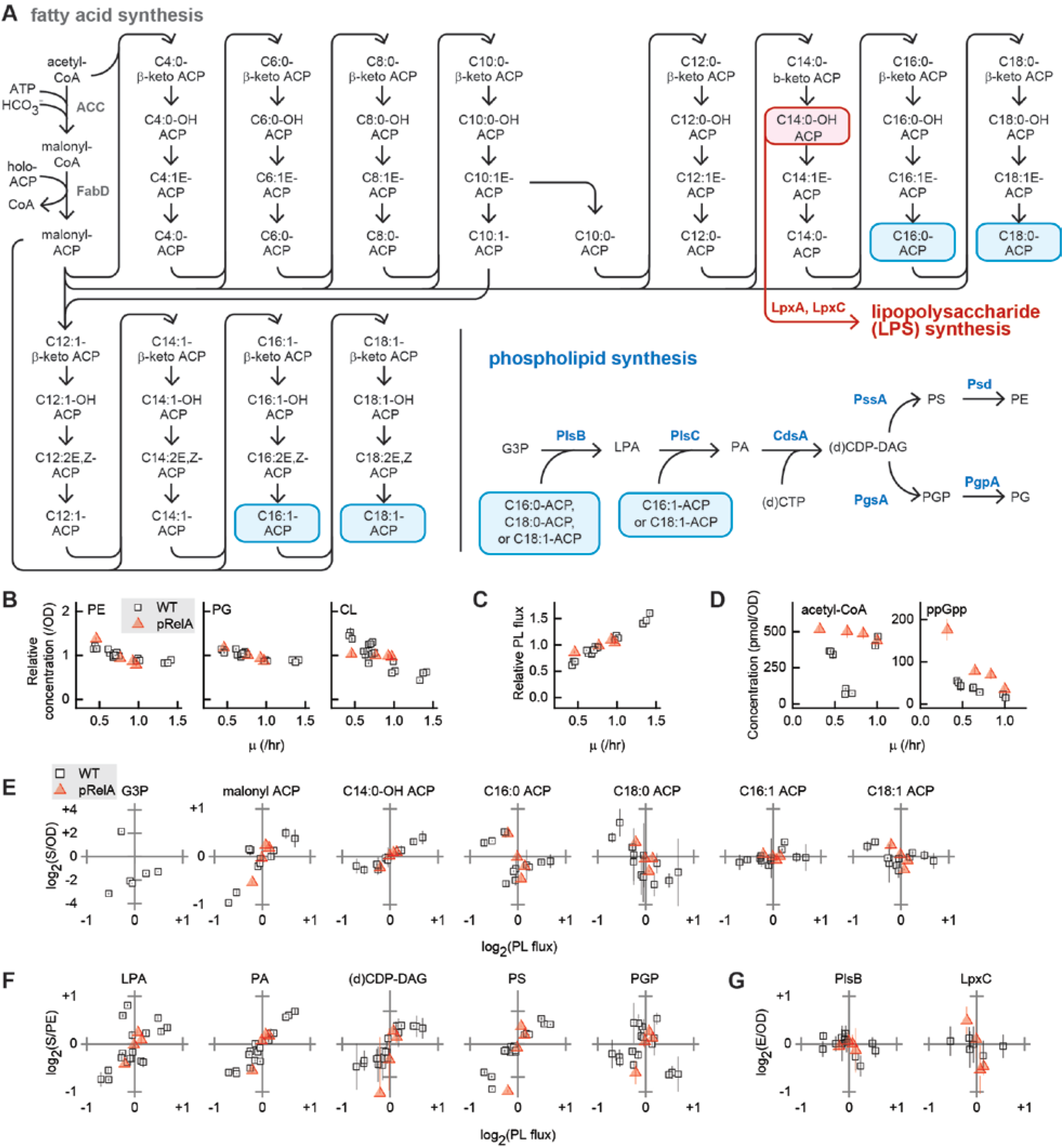
Characterization of the *E. coli* fatty acid and PL synthesis pathways during steady-state growth. **A.** The fatty acid and PL synthesis pathways. Long-chain fatty acids are transferred from C16:0-ACP, C18:0-ACP, or C18:1-ACP (highlighted in blue) to *sn-*glycerol-3-phosphate (G3P) by PlsB to yield lysophosphatidic acid (LPA), which is acylated with C16:1-ACP or C18:1-ACP to generate phosphatidic acid (PA). PA is converted to (d)CDP-diacylglycerol ((d)CDP-DAG) with either CTP or dCTP. At the branch point in PL synthesis, (d)CDP-DAG is converted to phosphatidylglycerol phosphate (PGP), which is dephosphorylated to yield PG (~25% of total PL). (d)CDP-DAG is also a precursor of phosphatidylserine (PS), which is decarboxylated to yield PE (~70% of total PL). PG is further converted to CL (~5% of total PL). C14:0-OH-ACP (highlighted in red) is a precursor for the synthesis of lipopolysaccharide (LPS), the major component of the outer leaflet of the outer membrane. **B, C.** Growth rate (μ)-dependent abundance of PG, PE, and CL per OD (**B**) and μ-dependence of PL flux (**C**), normalized to average values for each strain. **D.** Steady-state concentrations of acetyl-CoA and ppGpp. **E, F, G.** Correlations of G3P and acyl-ACP concentrations **(E)**, PL intermediate concentrations **(F)**, and PlsB and LpxC concentrations **(G)** with PL flux. Concentrations of soluble substrates and enzymes are calculated using cell volume (proportional to OD), while PL intermediate concentrations are calculated using membrane volume, which is proportional to total PE. All concentrations and fluxes are normalized and log(2)-transformed for comparison. All data points and error bars represent the average and standard deviation, respectively, of measurements of three samples from one culture. For WT PL and PL intermediates, three independent biological replicates of each condition are depicted; for ACP species and PlsB, two independent biological replicates; for nucleotides, LpxC, and G3P, data from one experiment per condition is shown. For pRelA, one measurement per inducer concentration is shown.

### Growth regulates PL flux via post-translational control of PlsB

We first sought to identify the control points of PL synthesis by determining correlations between PL flux and concentrations of pathway intermediates during steady-state growth. If PL and fatty acid synthesis rates are controlled by substrate concentrations, a positive correlation between substrate concentrations and flux should be observed. We quantified nucleotides, G3P, acyl-ACP, and PL synthesis intermediates using LC/MS (19). No correlation was observed between concentrations of the fatty acid precursor acetyl-CoA and μ (**Figure 1D**). We also found no significant positive correlation between PL flux and PlsB substrates C16:0-ACP or C18:1-ACP (Pearson correlation coefficients *r* <0.4, two-tailed test of significance *p* >0.1), while C18:0-ACP concentrations correlate negatively with PL flux (*r* = −0.76, *p* <0.01, **Figure 1E, Table 1).** Although G3P concentrations correlate with PL flux (*r* >0.99, *p* <0.01), growth in glycerol medium increases G3P by 20-fold without increasing PL abundance or flux. The absence of any significant positive correlation between ACP substrate concentrations and PL flux indicates that steady-state PL synthesis is not controlled by substrate concentrations.

**Table 1.** Pearson correlation coefficients and significance determined for PL flux and species abundance during steady-state growth in wild-type *E. coli. N* indicates the number of independent measurements used to calculate the correlation coefficients. Shaded cells indicate *p* < 0.05.

| Substrate | N | Pearson correlation coefficient (*r*) | Significance (*p*, two-tailed) |
| --- | --- | --- | --- |
| malonyl-ACP | 12 | 0.80 | <0.01 |
| G3P* | 5 | 0.99 | <0.01 |
| C14:0-OH-ACP | 12 | 0.94 | <10<sup>-4</sup> |
| C14:0-ACP | 12 | -0.16 | 0.6 |
| C16:0-ACP | 12 | -0.42 | 0.2 |
| C18:0-ACP | 12 | -0.76 | <0.01 |
| C16:1-ACP | 12 | 0.42 | 0.2 |
| C18:1-ACP | 12 | -0.38 | 0.2 |
| LPA | 18 | 0.68 | <0.01 |
| PA | 18 | 0.95 | <10<sup>-4</sup> |
| (d)CDP-DAG | 18 | 0.89 | <10<sup>-4</sup> |
| PS | 18 | 0.91 | <10<sup>-4</sup> |
| PGP** | 15 | 0.61 | 0.015 |

\*G3P: calculated with measurement in glycerol medium excluded.
\*\*PGP: calculated with measurements in glucose + cas-amino acids excluded.

**Table 1.** Pearson correlation coefficients and significance determined for PL flux and species 2 abundance during steady-state growth in wild-type *E. coli*. *N* indicates the number of 3 independent measurements used to calculate the correlation coefficients. Shaded cells indicate 4 $p < 0.05$ .
| Enzyme | N | Pearson correlation coefficient (*r*) | Significance (*p*, two-tailed) |
| --- | --- | --- | --- |
| LpxC | 6 | -0.4 | 0.4 |
| PlsB | 12 | -0.4 | 0.2 |
| PlsC | 12 | 0.63 | 0.03 |
| CdsA | 12 | 0.90 | <10<sup>-4</sup> |
| PssA | 12 | -0.4 | 0.2 |
| Psd | 12 | 0.2 | 0.6 |
| PgsA | 12 | -0.7 | 0.01 |
| PgpA | 12 | 0.9 | <10<sup>-4</sup> |
| AccA | 12 | 0.70 | 0.01 |
| AccB | 12 | 0.38 | 0.2 |
| AccC | 12 | 0.50 | 0.1 |
| AccD | 12 | 0.21 | 0.5 |
| AcpP | 12 | 0.04 | 0.9 |
| FabA | 12 | 0.29 | 0.4 |
| FabB | 12 | 0.28 | 0.4 |
| FabD | 12 | 0.35 | 0.3 |
| FabF | 12 | 0.76 | <0.01 |
| FabG | 12 | 0.68 | 0.02 |
| FabH | 6 | 0.49 | 0.3 |
| FabI | 12 | 0.82 | <0.001 |
| FabZ | 12 | 0.77 | <0.01 |
| GpsA | 12 | -0.69 | 0.01 |

In contrast to PL precursors, concentrations of the fatty acid initiation and elongation substrate malonyl-ACP correlate with PL flux **(Figure 1E,** *r* = 0.80, *p* <0.01, **Table 1)**, while the concentrations of most saturated ACP species remain relatively constant **(Supplemental Figure 1)** suggesting that the fatty acid initiation and elongation reactions are controlled primarily by malonyl-ACP concentrations and thus by the rate of malonyl-ACP synthesis by ACC and FabD. Concentrations of C14:0-OH-ACP, the lipid precursor of LPS, also correlate with PL flux (*r* = 0.94, *p* <10^-4^), suggesting that PL synthesis is naturally coupled with LPS synthesis via concentrations of C14:0-OH-ACP **(Figure 1E)**.

As with substrates, positive correlations between enzyme concentrations and reaction flux may indicate that a reaction rate is controlled by enzyme concentrations, i.e. via transcriptional control. We measured concentrations of fatty acid synthesis pathway enzymes using LC/MS to determine whether fatty acid flux is coupled with μ by transcription-based regulation. While concentrations of one of the four ACC subunits (AccA) and several of the fatty acid synthesis pathway enzymes correlate with PL flux **(Table 1, Supplemental Figure 2),** the increases are small (30%) relative to the PL flux increase (2-fold). We infer that malonyl-ACP synthesis and thus the rate of fatty acid synthesis are determined primarily by post-translational control of ACC. The negative correlation between C18:0-ACP and PL flux is consistent with long-chain acyl-ACP inhibiting malonyl-CoA synthesis by ACC via negative feedback inhibition (20).

Unlike long-chain acyl-ACP, the concentrations of all PL synthesis intermediates downstream of PlsB correlate positively with PL flux **(Figure 1F,** *r* = 0.61-0.99, **Table 1, Supplemental Figure 3**). The close match between the increase in concentration and PL flux is consistent with the rates of these reactions being under substrate control. The contrast between the trends in concentrations of the substrates and the products of PlsB implies that PlsB activity alone controls PL synthesis. PlsB activity might be regulated either via transcriptional control of PlsB concentration or via post-translational control of PlsB activity. Transcription of *plsB* is controlled by the membrane stress-activated sigma factor RpoE (21) and by the μ-sensitive regulator guanosine tetraphosphate (ppGpp) (22). Concentrations of ppGpp are inversely correlated with μ **(Figure 1D)**, implying that PL flux may be coupled to μ by transcriptional control of the *plsB* gene by ppGpp. However, we found no correlation between PL flux and PlsB concentration (**Table 1,** *p* >0.1). Instead, PlsB concentrations are nearly constant across all conditions studied **(Figure 1G)**. The invariance of PlsB concentrations over a 2-fold range of PL flux indicates that PlsB activity is regulated mainly via post-translational control. Unexpectedly, concentrations of LpxC, which catalyses the committed step in the LPS pathway (23), also do not correlate with PL flux (**Table 1,** *p >*0.1) and also remain stable **(Figure 1G),** indicating that flux into the LPS pathway is not varied between steady-state growth conditions by changes in LpxC concentration.

μ can be modulated independently of nutrient conditions by artificially varying ppGpp. Titrating ppGpp above basal concentrations, but below concentrations occurring during starvation, reduces steady-state μ by reducing ribosomal RNA synthesis (24). Furthermore, PlsB activity is inhibited at high ppGpp concentrations observed during amino acid starvation (25). We titrated μ by expressing the catalytic domain of the ppGpp synthesis enzyme RelA (RelA*) from an inducible promoter (PTet) in glucose medium. As previously observed (24), RelA*-titrated cultures exhibit ppGpp concentrations that are elevated 2-fold above wild-type cultures growing at similar rates **(Figure 1D).** Trends in PL abundance, PL flux, acyl-ACP species, PL intermediates, and PL synthesis enzyme concentrations in ppGpp-limited cultures closely follow the trends observed when μ is varied by carbon source **(Figure 1B-C, 1E-G, Supplemental Figures 1, 2, 3).** Notably, despite ppGpp regulation of *plsB* transcription, concentrations of PlsB also do not substantially vary with increasing concentrations of steady-state ppGpp. Interestingly, LpxC concentrations decrease with increasing PL flux in these conditions **(Figure 1G)**. The trends in substrate, enzyme, and intermediate concentrations observed in ppGpp-limited cultures are consistent with growth regulating PL flux via post-translational control – and not transcriptional control – of PlsB.

### Mathematical modelling supports PlsB control over steady-state PL and LPS synthesis

To test whether regulation of PlsB activity is sufficient to control steady-state PL synthesis, we constructed a simplified differential equation model that describes fatty acid, LPS initiation, and PL biosynthesis **(Figure 2A).** The model includes competitive inhibition of ACC by both C16:0-ACP and C18:0-ACP as the sole regulatory interactions. The model also includes a branch point at C14:0-OH-ACP into LPS synthesis. LPS initiation is represented in the model by a single step that combines reactions catalysed by LpxA and LpxC, as LpxC catalyses the first irreversible reaction in the LPS pathway. Concentrations of G3P and C16:1-ACP are fixed in the model to reflect experimental observations of PlsB saturation by G3P and invariance of C16:1-ACP concentrations, respectively (see **Supplementary Methods** for model details, parameter sets, and sensitivity analysis).

**Figure 2.**
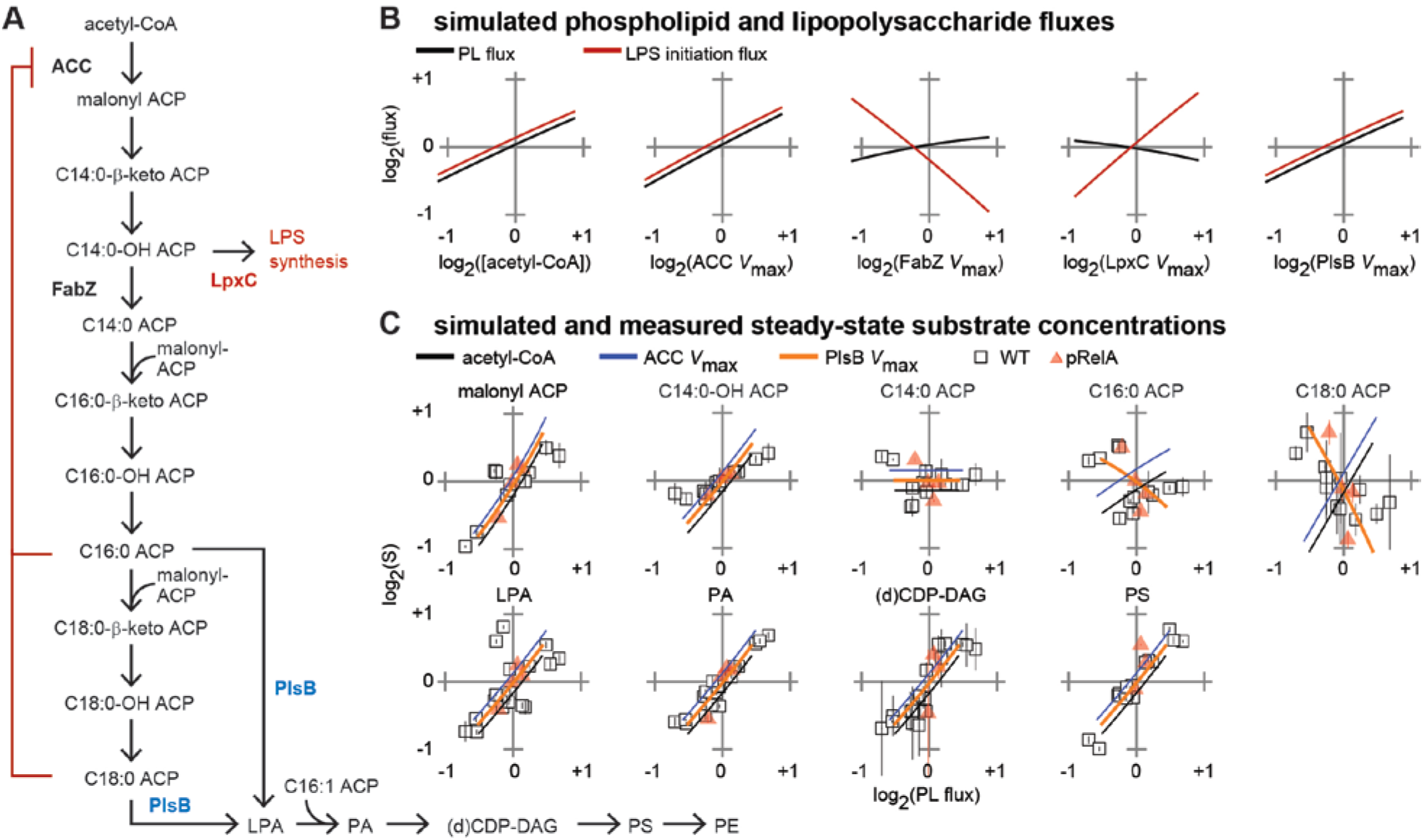
Simulated PL synthesis and experimental data identifies PlsB as site of PL synthesis control during steady-state growth. **A.** Reactions simulated in the model. For simplicity, only reactions late in the saturated fatty acid pathway are included, and branching of the PL pathway into PE and PG is not included. Each reaction is modelled as an irreversible one- or two-substrate Michaelis-Menten reaction. ACC is competitively inhibited by C16:0-ACP and C18:0-ACP. Reactions catalysed by FabI/FabZ and LpxA/LpxC are considered in the model as single reactions represented in the diagram by “FabZ” and “LpxC”, respectively. **B.** Response of PE and LPS fluxes to varying *V*_max_ of pathway enzymes and acetyl-CoA concentrations. 4-fold variations in *V*_max_ of all other reactions tested did not change PE or LPS flux (not shown). **C.** Simulated changes in metabolite concentrations in response to variations in PlsB and ACC *V*_max_ and acetyl-CoA (line plots) overlaid on experimentally-measured concentrations (scatter plots, data from **Figure 1**). Line plots are offset to prevent overlap. Differential equations and parameters of the simulation are provided in the **Supplemental Methods**.

We used the model to identify enzymes and substrates that exert control over PL and LPS synthesis. Increasing either ACC or PlsB *V*_max_ by 4-fold increased simulated PL and LPS fluxes in parallel by 2-fold **(Figure 2B)**, while changing *V*_max_ of all other enzymes in the simulated pathway had little or no effect on simulated PL flux. The insensitivity of PL flux to *V*_max_ of all fatty acid and PL synthesis enzymes aside from ACC and PlsB suggests that the experimentally-observed positive correlations between enzyme concentrations and PL flux do not actually increase PL flux. The unexpectedly strong control of LPS flux by PlsB is due to the natural coupling between PL flux and concentrations of the LPS synthesis substrate C14:0-OH-ACP. Changes in the C14:0-OH-ACP dehydration/reduction *V*_max_ (catalysed in *E. coli* by FabZ and FabI, respectively) and LPS initiation *V*_max_ (catalysed by LpxA and LpxC) exert strong and opposing control over LPS flux. However, LpxC and FabZ do not exert significant control over PL synthesis, indicating that substantial flux can be diverted into the LPS pathway without depleting PL synthesis. Of the two substrates considered in the model (acetyl-CoA and C16:1-ACP), only acetyl-CoA affects PL and LPS flux **(Figure 2B).**

To determine which of the two enzymes with non-negligible PL flux control – ACC or PlsB – actually controls PL flux during steady-state growth, we compared the predicted trends in substrate concentrations caused by simulated *V*_max_ variations against experimentally-observed trends. The predicted trends driven by ACC and PlsB *V*_max_ variation closely reproduce most experimentally-observed trends in both fatty acid and PL intermediate concentrations **(Figure 2C)**. However, variations in ACC and PlsB *V*_max_ cause opposing trends in predicted concentrations of PlsB substrates C16:0-ACP and C18:0-ACP, allowing ACC- and PlsB-based regulation to be distinguished. Increasing ACC *V*_max_ causes C16:0-ACP and C18:0-ACP concentrations to increase with PL flux, which contradicts our experimental observations. However, increasing PlsB *V*_max_ is predicted to decrease C16:0-ACP and C18:0-ACP concentrations, a trend that better follows the experimentally-observed trends (**Figure 2C).** We therefore conclude that PL synthesis is primarily regulated by PlsB during steady-state growth. Our model also suggests that PlsB control over steady-state LPS flux (achieved by its tight control over C14:0-OH-ACP concentrations) may be sufficient to couple LPS synthesis with growth as well.

### Translation inhibition causes carbon overflow into fatty acid synthesis

We set out to evaluate whether a known regulator of PL synthesis, ppGpp, is able to directly regulate PlsB activity during steady-state growth. High concentrations of ppGpp lead to PlsB inhibition *in vivo* (25), although it remains unclear whether this inhibition acts via a ppGpp-PlsB interaction or via an indirect route. Low concentrations of ppGpp correlate inversely with μ **(Figure 1D).** The notion that ppGpp might directly control PL synthesis even at the low basal concentrations present during steady-state growth has been proposed but never tested (26). Acyl-ACP substrates of PlsB increase as ppGpp is titrated upwards with RelA* while the product of PlsB (LPA) decreases, consistent with ppGpp inhibiting flux into the PL pathway (**Figure 1E, 1F**), and closely following simulated variations in PlsB *V*_max_ **(Figure 2C).** Therefore, changes in μ driven by sub-stringent concentrations of ppGpp must somehow influence steady-state PlsB activity. However, the mode of control by ppGpp (transcriptional control of *plsB*, post-translational inhibition of PlsB, or indirect control via an unknown regulator of PlsB) cannot be determined from steady-state data alone.

We first wished to observe the effects of high concentrations of ppGpp on the fatty acid and PL synthesis pathways. Synthesis of high concentrations of ppGpp by RelA (the stringent response) is triggered by any stress or starvation conditions that lead to the specific biochemical cue of uncharged tRNA bound to the ribosome. During the stringent response, ppGpp accumulates by more than 10-fold over basal concentrations and PL synthesis rates are reduced by approximately half (25, 26), likely due to PlsB inhibition (27). We triggered ppGpp synthesis by adding the tRNA aminoacylation inhibitor mupirocin to glucose cultures of wild-type *E. coli*. To distinguish effects of ppGpp from effects of translation inhibition, mupirocin was also added to a culture of Δ*relA E. coli*. Mupirocin causes a 10-fold accumulation of ppGpp in the wild-type strain within 1 minute to over 400 pmol/OD, reaching >800 pmol/OD after 3 minutes **(Supplemental Figure 4).** Unexpectedly, mupirocin treatment also transiently increased malonyl-ACP and all hydroxyl-ACP species at the expense of holo-ACP in both strains **(Figure 3A, Supplemental Figure 5)**. The increases in malonyl- and hydroxyl-ACP concentrations are matched by a corresponding increase in PL synthesis intermediates, suggesting that mupirocin diverts a pulse of carbon into the fatty acid and PL synthesis pathways **(Figure 3B, Supplemental Figure 6)**. In the wild-type strain, the pulse of fatty acid synthesis activity is rapidly followed by C14:0-, C16:0-, and C18:0-ACP accumulation, consistent with ppGpp inhibition of PlsB **(Figure 3A, Supplemental Figure 5)**. C16:0-ACP accumulation is followed in turn by an increase in holo-ACP and a decrease in malonyl-ACP, consistent with ACC inhibition **(Figure 3A)**. PL intermediates PA, PS, and PGP are rapidly depleted in the wild-type strain after briefly increasing **(Figure 3B).** The transient carbon influx briefly shifts the population of the acyl chains incorporated into PL towards longer-chain fatty acids, likely due to increased malonyl-ACP concentrations favouring fatty acid elongation over membrane incorporation by PlsB and PlsC **(Supplemental Figure 6).** While suppression of fatty acid synthesis in the wild-type strain can be attributed to ppGpp, it is unclear what attenuates the carbon influx in the Δ*relA* strain; however C14:0-ACP accumulation after 10 minutes likely contributes to ACC inhibition. Inhibition of fatty acid elongation also depletes the LPS precursor C14:0-OH-ACP, which is expected to decrease LPS synthesis in parallel with PL synthesis **(Figure 3A)**.

**Figure 3.**
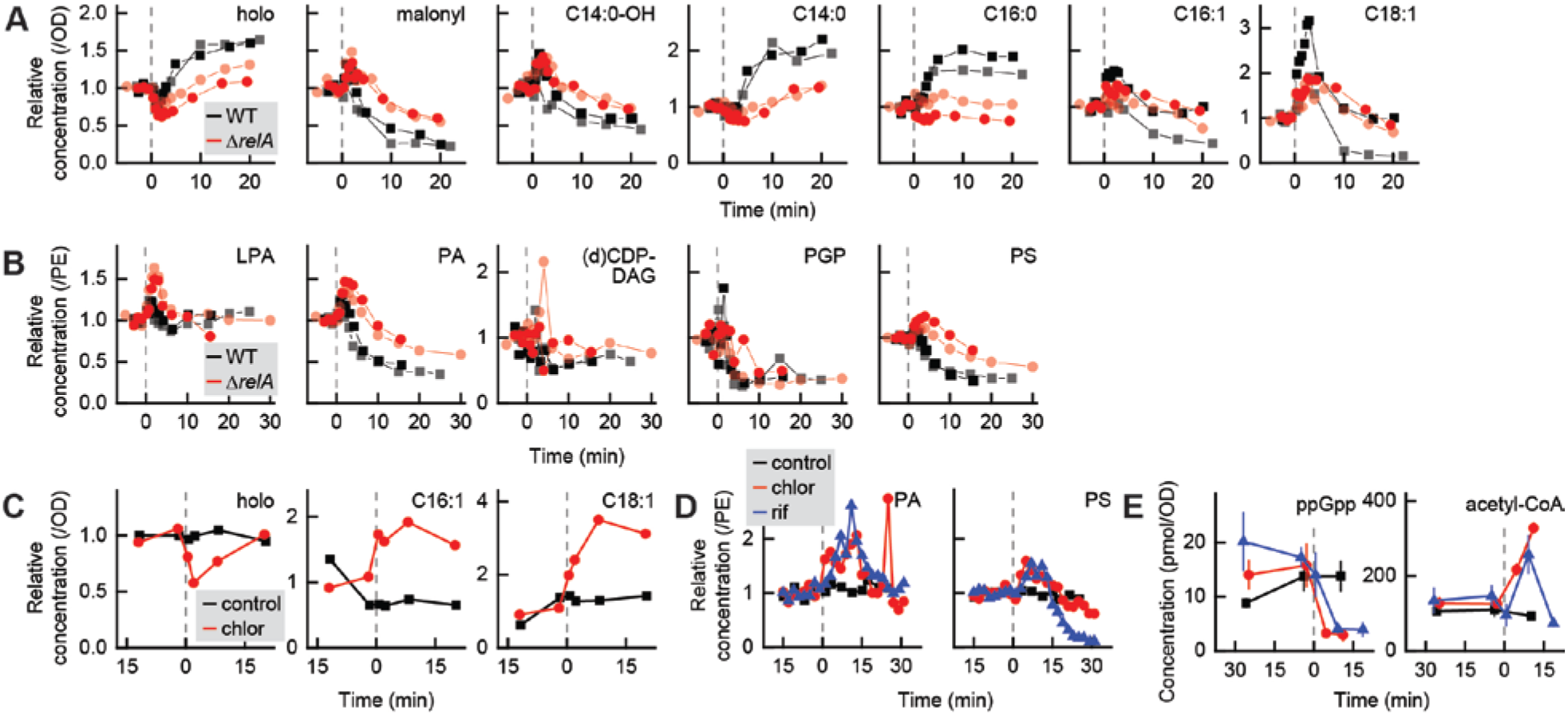
Responses of the fatty acid and PL synthesis pathways to translation inhibition. **A, B.** Responses of acyl-ACP **(A)** and PL intermediates **(B)** to mupirocin. Mupirocin was added at 0 min (indicated by dashed lines) to glucose cultures of *E. coli* wild-type and Δ*relA* NCM3722. Each point represents one measurement from a time series collected from an independent culture. Two biological replicate series for each strain are depicted. **C, D.** Addition of translation inhibitor chloramphenicol or the transcription inhibitor rifampicin to glycerol cultures of wild-type *E. coli* (indicated by dashed lines at 0 min) causes an influx of carbon into the fatty acid pathway, as suggested by a decrease in holo-ACP and an increase in unsaturated long-chain acyl-ACP species **(C).** The pulse of carbon continues into the PL synthesis pathway, as indicated by transient increases in total PA and PS **(D).** Each trajectory in **C-D** is obtained from a single culture. **E.** Chloramphenicol and rifampicin both trigger a rapid decrease in ppGpp and accumulation of ACC and FabH substrate acetyl-CoA. Points and error bars in **E** are averages and standard deviation of triplicate samples, respectively, from individual experiments.

As malonyl-ACP increases transiently after mupirocin in both wild-type and Δ*relA* strains, we hypothesized that translation inhibition somehow diverts carbon into lipid synthesis. We added the ribosome inhibitor chloramphenicol and the transcription initiation inhibitor rifampicin to glycerol cultures of wild-type *E. coli*. Both compounds inhibit translation via mechanisms that suppress ppGpp synthesis. As with mupirocin treatment of the Δ*relA* strain, chloramphenicol triggered a rapid decrease in holo-ACP and an increase in long-chain unsaturated acyl-ACP species C16:1-ACP and C18:1-ACP **(Figure 3C).** Both antibiotics triggered an increase in PL synthesis intermediates PA and PS that resembled the response of the Δ*relA* strain to mupirocin **(Figure 3D).**

What might cause the transient increase in fatty acid synthesis observed after translation inhibition? Interestingly, both rifampicin and chloramphenicol increased acetyl-CoA concentrations in glycerol cultures by 3-fold **(Figure 3E)**, suggesting a possible cause. Acetyl-CoA also increased in both wild-type and Δ*relA* strains after mupirocin treatment in glucose medium, before decreasing ~30% in the wild-type strain **(Supplemental Figure 4).** While it is unclear why translation inhibition would increase acetyl-CoA, the observed response of the fatty acid pathway is consistent with our mathematical model, which predicts fatty acid flux to be highly sensitive to changes in acetyl-CoA concentrations **(Figure 2B)**. We confirmed the sensitivity of the fatty acid and PL synthesis pathways to environmental changes using a fast nutritional upshift. Addition of glucose and amino acids to a glycerol culture also caused rapid accumulation of PA and PS species that resemble the increases observed following translation inhibition **(Supplemental Figure 7).**

### Moderate to high concentrations of ppGpp regulate PL synthesis via post-translational control

In order to clearly discern the effects of ppGpp on the fatty acid and PL synthesis pathways without complications introduced by translation inhibition, we monitored fatty acid and PL synthesis pathways immediately after inducing RelA*. The responses of the fatty acid and PL synthesis pathways are again consistent with PlsB inhibition causing long-chain acyl-ACP to accumulate, which depletes malonyl-ACP by inhibiting ACC **(Figure 4A)**. PL intermediates also respond in a manner consistent with PlsB inhibition: LPA species steadily decrease, followed by PA, PS, and PGP **(Figure 4B)**. Addition of chloramphenicol 10 minutes following RelA* induction causes an increase in unsaturated long-chain acyl-ACP, though ppGpp appears to attenuate the response of the PL pathway to translation inhibition.

**Figure 4.**
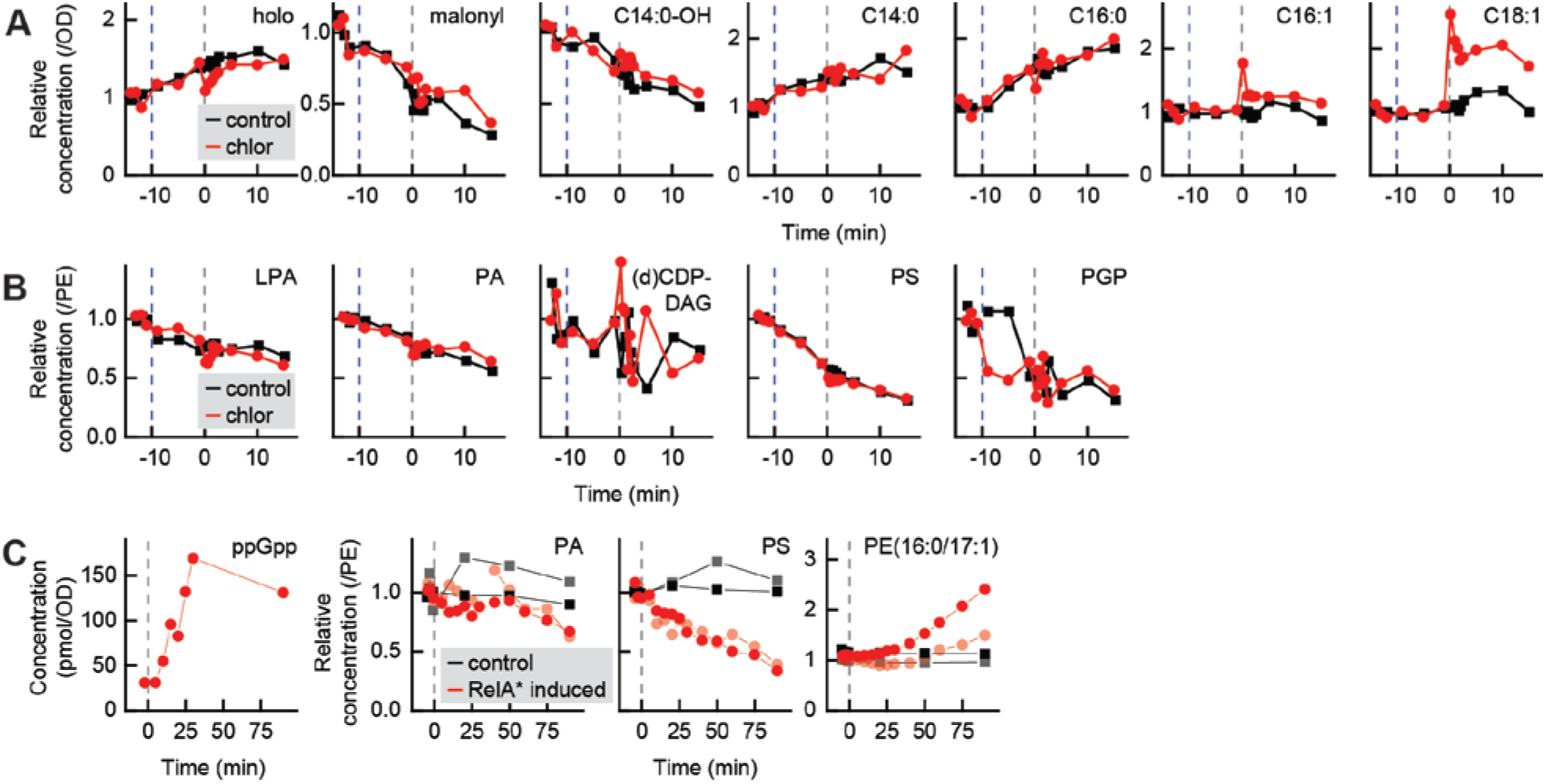
PlsB activity is suppressed by moderate to high concentrations of ppGpp via post-translational inhibition. **A, B.** Response of acyl-ACP **(A)** and PL intermediate pools **(B)** to maximal overexpression of ppGpp synthesis enzyme RelA*. RelA* expression was induced by addition of 40 ng/mL doxycycline at −10 minutes (dashed blue line), followed 10 minutes later by chloramphenicol addition (dashed grey line at 0 min). **C.** Response of concentrations of PL intermediates and cyclopropyl-PE to mild RelA* overexpression, triggered by addition of 1 ng/mL doxycycline (dashed grey line at 0 min). Each time series represents one independent biological replicate.

We next sought to evaluate whether ppGpp concentrations far below those occurring during the stringent response are also capable of regulating PlsB via post-translational control. As the effects of post-translational control can be distinguished from transcriptional regulation by response time, we closely followed the dynamics of the PL pathway after mildly inducing ppGpp synthesis with a low doxycycline concentration. ppGpp increased within 10 minutes of RelA* induction and reached its elevated steady-state concentration (~150 pmol/OD) by 20 minutes. Concentrations of PS begin to decrease within 15 minutes (Figure 4C), consistent with post-translational inhibition of PL synthesis. To verify that the immediate PS depletion is too fast to be explained by transcriptional regulation, we compared the PS response kinetics with the kinetics of a ppGpp-driven response known to be mediated by transcriptional control. Cyclopropyl PL are produced from unsaturated fatty acids of membrane PL by the enzyme Cfa, expression of which is induced via transcriptional control by ppGpp (28). Cyclopropyl PL begins to accumulate 25 minutes after RelA* induction, well after the PS decrease is established. We conclude that ppGpp concentrations well below stringent response concentrations inhibit PlsB via a post-translational mechanism.

## Discussion

Over several decades, biochemical and genetic research has extensively characterized the pathways and the enzymes that produce the building blocks of bacterial membranes (4, 29, 30). Many control mechanisms acting on both transcriptional and post-translational levels have been proposed to maintain membrane homeostasis. For instance, each of the enzymes ACC (31), FabH (32), FabZ (33), FabI (34), LpxC (35), and PlsB (36) have been suggested to contribute to membrane synthesis regulation. However, the existence of a control mechanism does not necessarily indicate that it is used to regulate total membrane synthesis flux during steady-state growth. Furthermore, models derived from biochemical and genetic studies can overlook self-regulating mechanisms that occur automatically in a metabolic pathway, such as the coupling of C14:0-OH-ACP concentrations to PL flux. The primary value of our work is that it reveals the regulatory mechanisms that are actually used by growing *E. coli* to regulate membrane biosynthesis. Both our data and our modelling indicate that only two of the many control mechanisms previously identified are sufficient to regulate total PL biosynthesis during steady-state growth. Specifically, we find that cellular demand for membrane synthesis is communicated to PlsB via an undiscovered post-translational mechanism. In turn, PlsB activity regulates fatty acid synthesis by consuming long-chain acyl-ACP species, relieving feedback inhibition of ACC (20) and increasing fatty acid flux. This demand-level flux regulation is consistent with metabolic control theory (37).

Our measurements of enzyme concentrations indicate that PL flux is regulated primarily via post-translational control of enzyme activity, rather than via transcriptional regulation. Concentrations of pathway enzymes with the greatest control over PL flux (ACC and PlsB) are maintained at nearly invariant levels across a 3-fold range of μ. Although it has been long accepted that PlsB activity is adjusted via post-translational control and not transcriptional control (4), the strong flux control of ACC (31) and the growth-regulated transcription of the *accBC, accA*, and *accD* genes (38) inspired the suggestion that PL flux might be coupled with μ via transcriptional regulation. Instead, we find that transcriptional regulation of fatty acid and PL synthesis genes stabilizes the concentrations of fatty acid and PL synthesis enzymes across μ. The correlations between concentrations of several enzymes and PL flux that we observe (e.g. FabF) are unlikely to increase total PL flux, but rather regulate aspects of membrane metabolism aside from PL flux, such as membrane fluidity. Maintaining a stable membrane synthesis capacity enables the cell to quickly respond to any change in membrane demand at the cost of expressing enzymes that remain less active at slow or moderate μ (39).

Our data are also essential for evaluating proposed models of LPS synthesis regulation. Most importantly, both our model and our experiments indicate that PL flux – and thus, PlsB activity – controls concentrations of LPS precursor C14:0-OH-ACP. This tightly links fluxes into the PL and LPS synthesis pathways, as varying PL synthesis by adjusting either PlsB or ACC activity will naturally vary LPS synthesis in parallel. Surprisingly, we find that flux into the LPS synthesis pathway is not adjusted by variations in LpxC concentrations, as LpxC concentrations remain constant despite a 3-fold change in μ. LPS flux may be varied instead by FabZ, concentrations of which increase by 50% over the 3-fold range of μ sampled **(Supplemental Figure 2)**. Our model predicts this would attenuate the effects of increasing C14:0-OH-ACP concentrations caused by elevated fatty acid synthesis and divert flux away from LPS synthesis without affecting PL flux. LpxC concentrations may be stabilized across μ by degradation by the FtsH protease (40). Degradation of LpxC by FtsH might be used as an emergency brake to rapidly halt LPS synthesis during growth arrest or growth transitions to prevent accumulation of toxic LPS intermediates. A comprehensive characterization and modelling (41) of LPS substrates and enzymes across several steady-state conditions, as we have done for PL synthesis, will be necessary to determine how LPS synthesis is varied.

Although PlsB determines PL flux during steady-state growth, this control is not absolute, as fatty acid and PL flux (and likely LPS flux) are also sensitive to acetyl-CoA concentrations and ACC activity. The sensitivity of membrane synthesis to protein synthesis arrest demonstrates the responsiveness of membrane synthesis to carbon metabolism. While this sensitivity facilitates rapid adaptation of PL flux to environmental changes, the tight connection between protein synthesis, central carbon metabolism, and membrane biogenesis demands additional regulation (including inhibition by ppGpp) to re-balance the pathway with μ and prevent PL overflow. If protein synthesis is inhibited in a manner that does not decrease carbon flow into the fatty acid pathway (e.g. by nitrogen starvation), continued PL and LPS synthesis would outpace synthesis of the lipoproteins that tether the outer membrane to the peptidoglycan layer. Inhibition of PL and LPS pathways by ppGpp would thus prevent production of excess membrane, enforcing the coupling of PL and lipoprotein synthesis. Consistent with this notion, Δ*relA* strains generate higher quantities of extracellular PL and LPS, likely as outer membrane vesicles (42).

Identifying the specific signal that controls PlsB will reveal how membrane synthesis is synchronized with growth. But what controls PlsB? Post-translational control of PlsB by moderately-high concentrations of ppGpp (≥150 pmol/OD) at least partly contributes to steady-state membrane synthesis regulation, but likely only during nutritional downshifts, stress, or very slow steady-state growth (μ ≪ 0.5 hr^-1^) when such concentrations are achieved. Whether basal ppGpp concentrations observed during fast growth (μ > 0.5 hr^-1^, ppGpp < 60 pmol/OD) also regulates PL synthesis via PlsB control remains unclear. *In vivo* experiments such as ours cannot determine the mechanism by which ppGpp inhibits PlsB. Although it has been demonstrated that high concentrations of ppGpp directly inhibits PlsB (25), multiple groups have been unable to observe ppGpp inhibition of PlsB *in vitro* when acyl-ACP are used as substrates (43, 44). ppGpp may therefore inhibit PlsB indirectly via a regulator or a cellular process that interacts with PlsB. Immunoprecipitation experiments found several proteins that interact with PlsB including ACP and PssA, as well as several whose roles are unclear (PlsX and YbgC) (45), suggesting that PlsB forms part of a PL synthesis complex that may couple PlsB activity with μ. As PL synthesis flux has been observed to oscillate with the cell division cycle in *E. coli* and other bacteria (46), PlsB may also be regulated by the divisome or by septum formation. Degradation of PL by phospholipases may also play an important role in membrane homeostasis during steady-state growth (47), as is known to occur during growth and division in eukaryotes (48). In addition to identifying the allosteric regulators of PlsB, studies that integrate connections between PL catabolism and transport with PL synthesis are needed for a comprehensive understanding of membrane homeostasis.

## Acknowledgements

We thank Michael Cashel for providing *E. coli* strain CF7974. We thank Christophe Danelon, Bertus Beaumont, and Frank Bruggeman for invaluable suggestions for improving the manuscript. We thank the National BioResource Project (NIG, Japan) for sharing the LpxC overexpression plasmid. This research was funded by a grant to GB from the Netherlands Organisation for Scientific Research (ALW Open 824.15.018) and an award from the Frontiers of Nanoscience program.

## Author contributions

MN and GB designed the experiments. MN, FB, NvdB, NS, FY, and GB performed experiments. MN, NvdB, and NI developed analytical methods. MN, FY, and GB analysed the data. GB supervised the research, constructed the mathematical model, and wrote the manuscript.

## Materials and Methods

### Culture conditions

Cultures were grown in 250-1000 mL Erlenmeyer flasks filled up to 10% of nominal volume with MOPS minimal medium (49) with 9.5 mM NH_4_Cl or ^15^NH_4_Cl and 0.2% (w/v) carbon source (acetate, succinate, malate, glycerol, glucose, U-^13^C-glucose (Cambridge Isotope Laboratories) or glucose supplemented with 0.1% Cas-amino acids). Culture flasks were placed in a Grant Instruments Sub Aqua Pro dual water bath at 37 °C and agitated by stirring with a 12 mm magnetic stir bar (VWR), coupled to a magnetic stir plate (2mag MIXdrive 1 Eco and MIXcontrol 20) set at 1200 rpm. Growth was monitored by optical density measurement at 600 nm using Ultrospec 10 Cell Density Meter (GE Healthcare). Samples for acyl-ACP, lipid analysis and proteomics were collected using cultures without isotopic labelling. Samples for nucleotide phosphate measurements were collected from U-^15^N-labeled cultures and samples for G3P measurements were collected from U-^13^C-labeled cultures.

### Strains and plasmid pRelA*

All experiments were performed using *Escherichia coli* K-12 strain NCM3722 (CGSC# 12355) and its derivatives. NCM3722 *relA::kan* was constructed by P1 phage transduction using strain CF7974 (MG1655 Δ*lac* (*rph*^+^) *relA255::kan*) as a donor. Plasmid pRelA* was created by cloning DNA encoding residues 1-455 from the *E. coli* RelA protein into BglBrick plasmid pBbS2k (50) (SC101* origin of replication, P_Tet_ promoter, kanamycin resistance). The fluorescent protein mVenus was fused by restriction-digestion to the C-terminus of RelA via a glycine-serine linker.

### Metabolite sampling

Samples for acyl-ACP, proteomics and lipid analysis were acquired by fast quenching of 1 mL of culture sample into 250 μL of ice-cold 10% trichloroacetic acid. After 10 min incubation at 0°C cells were pelleted by centrifugation and stored at −80°C until analysis. For nucleotide phosphates and polar metabolites analysis, samples were acquired by a modified fast vacuum filtration method (51). 1 mL of culture was collected by vacuum on a pre-wetted 2.5 cm 0.45 μm HV Durapore membrane filter. After rapid collection the filter was immediately placed upside-down in quenching solution. For the measurement of nucleotide phosphates, 1 mL of ice-cold 2 M formic acid with 10 μL internal standards mix was used as a quenching solution, which was subsequently neutralized by 25 μL of 28% ammonium hydroxide. For G3P measurements, 1 mL of 50:30:20 (v/v/v) mixture of methanol, acetonitrile and water with 0.1% formic acid with 10 μL internal standard solution (cooled on dry ice) was used as a quenching solution. After 10 min incubation cells were washed from the filter, transferred to a tube and stored at −80°C until analysis.

### Preparation of internal standards

Isotopically-labeled internal standards (IS) were used to control for sampling and measurement variation. For acyl-ACP and proteomics assays U-^15^N *E. coli* whole cell extracts were prepared using a MOPS minimal medium culture with ^15^NH_4_Cl as the sole nitrogen source. At OD of ~0.5 10% TCA was added 1:4 to the culture to facilitate quenching of metabolism. After 10 min incubation on ice 10 mL single-use IS aliquots were collected by centrifugation and stored at −80°C until the sample preparation. For the phospholipid measurement U-^13^C lipid extract was prepared using a culture grown in minimal MOPS medium with 0.2% U-^13^C glucose as the sole carbon source. At OD of ~0.5 10% TCA was added 1:4 to the culture and insoluble cell material was collected by centrifugation after 10 min incubation on ice. Pellets were resuspended in mixture consisting of 75 μL MeOH, 10 μL 15 mM citric acid/ 20 mM dipotassium phosphate buffer and 250 μL of methyl-t-butyl ether per 1 mL of initial culture volume. After vortexing and 10 min sonication phase separation was induced by addition of 70 μL/1mL of 15 mM citric acid/ 20 mM dipotassium phosphate buffer. After further vortexing, sonication and 10 minutes of incubation at room temperature phases were separated by 10 min centrifugation at 4000 rpm at room temperature. Upper phase was collected to a glass vial and stored at −20°C until sample preparation.

### Instrumentation

All LC/MS runs were performed using Agilent LCMS consisting of binary pump (G1312B), autosampler (G7167A), temperature-controlled column compartment (G1316A), and triple quadrupole (QQQ) mass spectrometer (G6460C) equipped with a standard ESI source, all operated using MassHunter data acquisition software (version 7.0). Mass spectrometer operated in dynamic MRM mode using transitions generated *in silico* by a script written in Python using RDkit library using chemical structures of target compound as input. Transitions for targeted proteomics assays were developed using Skyline (52) based on protein sequences from the Uniprot database.

### LC/MS quantification of acyl-ACP intermediates

Acyl-ACP were measured using a published method (19) with minor modifications. Lysis buffer was prepared by suspending appropriate number of frozen U-^15^N-labeled *E. coli* pellets in 10 mL of 50 mM potassium phosphate buffer, pH 7.2, 6 M urea, 10 mM N-ethyl-maleimide, 5 mM EDTA and 1 mM ascorbic acid. 1 mL of lysis buffer was added to each of TCA-quenched and pelleted cells and proteins were isolated by chloroform/methanol precipitation as described previously. Protein pellets were resuspended in 10 μL of digestion buffer (4% 2-octyl-glucoside in 25 mM potassium phosphate buffer, pH 7.2) and after adding 10 μL of 0.1 mg/mL GluC protease (Promega) incubated overnight at 37°C. After quenching by addition of 5 μL MeOH, samples were centrifuged and 10 μL was injected in LC/MS system. Separation was performed on 2.1 mm x 50 mm 1.7 μm CSH C-18 column (Waters) held at 80°C using a binary gradient: 15% B, 3 minute ramp to 25%, 9 min increase to 95% and 1 minute hold at 95% B before 3 minute re-equilibration at starting conditions (A: 25 mM formic acid, B: 50 mM formic acid) at a flow rate of 0.6 mL/min.

### LC/MS quantification of phospholipids

Phospholipids sample preparation procedure is a combination of an MTBE extraction method (53) and an established LC/MS method (54) Pelleted *E. coli* were resuspended in mixture containing 150 μL of MeOH, 250 μL of U-^13^C *E. coli* extract prepared as described above and 250 μL MTBE. After vigorous vortexing and sonication 125 μL of 15 mM citric acid/ 20 mM dipotassium phosphate buffer was added to homogenized pellets. Following further vortexing and 10 min incubation at room temperature, liquid phases were separated by centrifugation for 10 min at 20000g. 500 μL the of upper phase was moved to a new tube and dried in a vacuum centrifuge (Labconco). Dried lipid films were resuspended in 10 μL 65:30:5 (v/v/v) isopropanol/acetonitrile/H_2_O, supplemented with 10 mM acetylacetone. After resuspension, 5 μL H_2_O was added to reduce the organic content of the buffer and 5 μL of resulting mixture was injected into the LC/MS system. Separation was performed on 2.1 mm x 50 mm 1.7 μm CSH C-18 column (Waters) at 60°C with a flow rate of 0.6 mL/min using the following binary gradient: 25% B, ramp to 56%B in 6 min followed by linear increase to 80% B in 6 min, 2 min hold at 100% B and 3 min re-equilibration (A: 0.05% NH4OH in water, B: 0.05% NH4OH in 80% isopropanol 20% ACN).

### LC/MS quantification of nucleotides

Frozen cell extracts were defrosted by 1-2 minute incubation in a 37°C water bath and sonicated for 10 minutes in water ice slurry. After 10 minute centrifugation at 20000g, samples were loaded on 1 mL/30 mg Oasis Wax cartridge (Waters) preconditioned with 1 mL of MeOH and 1 mL 50 mM ammonium acetate buffer, pH 4.5. After washing with 1 mL ammonium acetate buffer, analytes were eluted with 200 μL of 2.8% ammonium hydroxide in MeOH:ACN:H_2_O 50:30:20 (v:v:v). After addition of 10 μL of 5% trehalose and brief vortexing samples were dried down in a vacuum centrifuge (Labconco). Dried trehalose-stabilized extracts were re-dissolved in 20 μL of MeOH:ACN:H_2_O 50:30:20 (v:v:v) and moved to an autosampler vial for analysis. Separation was performed on 2.1 mm x 100 mm 3.5 μm iHilic column (HILICON) or SeQuant Zic-cHILIC, 2.1mm x 100 mm, 3μm (Merck) at 0.3 mL/min using the following binary gradient: 100% B, ramp to 85%B in 1.5 min followed by 10 min isocratic hold at 85% B and linear decrease 30% B in 3 minutes with 2 minute hold at 30% B and 8 minute re-equilibration at initial conditions. (A: 3.75 mM ammonium acetate, 1.25 mM acetic acid, 2 mM acetylacetone in MQ, B: 11.25 mM ammonium acetate, 3.75 mM acetic acid, 2 mM acetylacetone in 80% ACN). Injection volume was 2 μL.

### LC/MS quantification of G3P

Stored metabolite extracts were dried down in a vacuum centrifuge (Labconco), re-dissolved in 20 μL of MeOH:ACN:H_2_O 50:30:20 (v:v:v) and moved to an autosampler vial for analysis. Separation was performed on 2.1 mm x 100 mm 3.5 μm iHilic column (HILICON) at 0.3 mL/min using the following binary gradient: 100% B, ramp to 80% B in 10 min followed by linear decrease to 30% B in 3 min, 2 min hold at 30% B and 8 min re-equilibration. Injection volume was 2 μL.

### LC/MS targeted protein quantification

Relative concentrations of enzymes was measured by targeted proteomics using a modified version of the acyl-ACP assay. Lysis buffer was prepared by suspending appropriate number of frozen U-^15^N-labeled *E. coli* pellets in 10 mL of 50 mM potassium phosphate buffer, pH 7.2 and 6 M urea. To improve the detection of peptides from PlsB and LpxC, U-^15^N-labeled *E. coli* NCM3722 strains overexpressing PlsB and LpxC were used as internal standards. LpxC was overexpressed using the corresponding plasmid from the ASKA library (55) (National BioResource Project (NIG, Japan)). 1 mL of lysis buffer was added to each of TCA-quenched and pelleted cells and proteins were isolated by chloroform/methanol precipitation as described previously. Protein pellets were resuspended in 10 μL of digestion buffer (4% 2-octyl-glucoside in 25 mM Tris buffer, pH 8.1 supplemented with 1 mM CaCl_2_ and 5 mM TCEP). Alkylation of cysteine residues was performed by adding 3 μL of 50 mM iodoacetamide followed by 15 minutes of incubation in darkness. Digestion was performed by adding 10 μL of 0.2 mg/mL Trypsin Gold (Promega) and overnight incubation at 37°C. Samples were centrifuged and 10 μL was injected in LCMS system. Separation was performed on 2.1 mm x 50 mm 1.7 μm CSH C-18 column (Waters) held at 40°C using a binary gradient: 2% B, 20 minute ramp to 25% B, 4 min increase to 40% B, 0.5 ramp to 80% and 1 minute hold at 80% B before 3 minute re-equilibration at starting conditions (A: 25 mL formic acid, B: 50 mM formic acid) at a flow rate of 0.5 mL/min.

### Data Analysis

All LC-MS data files were processed in Skyline versions 4.x using target list based on *in silico* generated transition list. Each target compound had matching isotopically-labeled internal standard (IS). Processed data were exported as target compounds and IS peak areas and processed further using a set of Python scripts. Growth rates were obtained from linear fits to log-transformed growth curves. OD-corrected data were obtained by dividing the signal by OD600 value interpolated from growth curve at the time of sampling. PE-corrected results were produced by dividing the signal by sum all of signals for all phosphatidyl-ethanolamine species from the same measurement (in case of phospholipids) or matching sample (in case of other assays). In nucleotide phosphate and G3P assay absolute concentrations were estimated based on amounts of internal standards in IS-spike solution assuming RR = 1 implies equimolar amounts of target compound and IS at the moment of quenching. Correlations in Table 1 were calculated from log(2) normalized concentrations and PL fluxes using the Descriptive Statistics function (OriginPro v. 2015).

### Mathematical modeling

The computational model was constructed and tested using COPASI version 4.24 (56). Full details of the model are described in the **Supplemental Methods**.

