## Supplemental Material for "Post-translational control of PlsB is sufficient to coordinate membrane synthesis with growth in *Escherichia coli*"

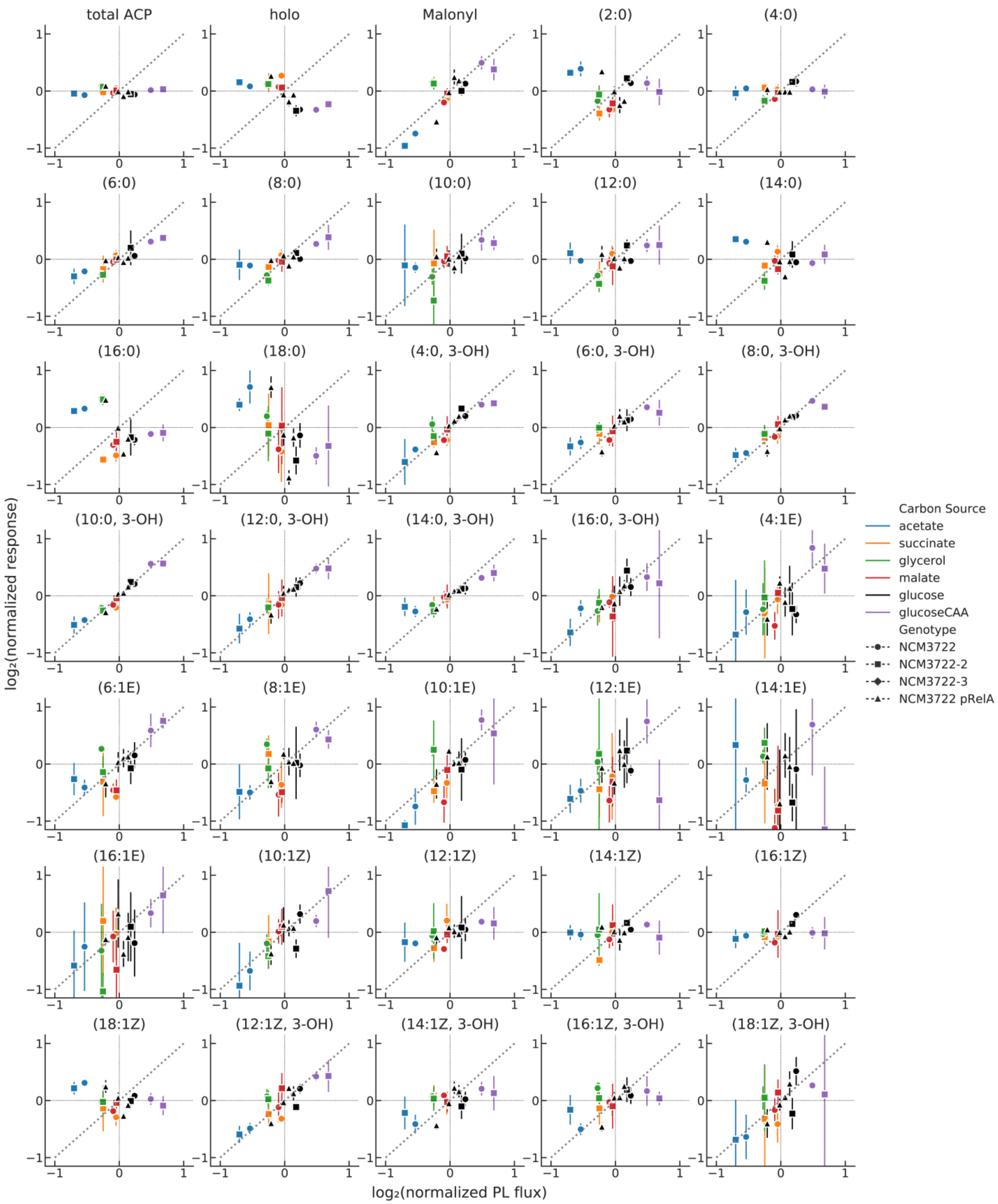

**Supplemental Figure 1.** Steady-state concentrations of acyl-ACP of *E. coli* NCM3722 in six media and of *E. coli* pRelA* in glucose cultures with titrated RelA* expression (triangles). Concentrations and PL fluxes are normalized to average across each individual replicate series. Error bars represent standard deviation of 3 sampling replicates. Two series of independent biological replicates were obtained for *E. coli* NCM3722.

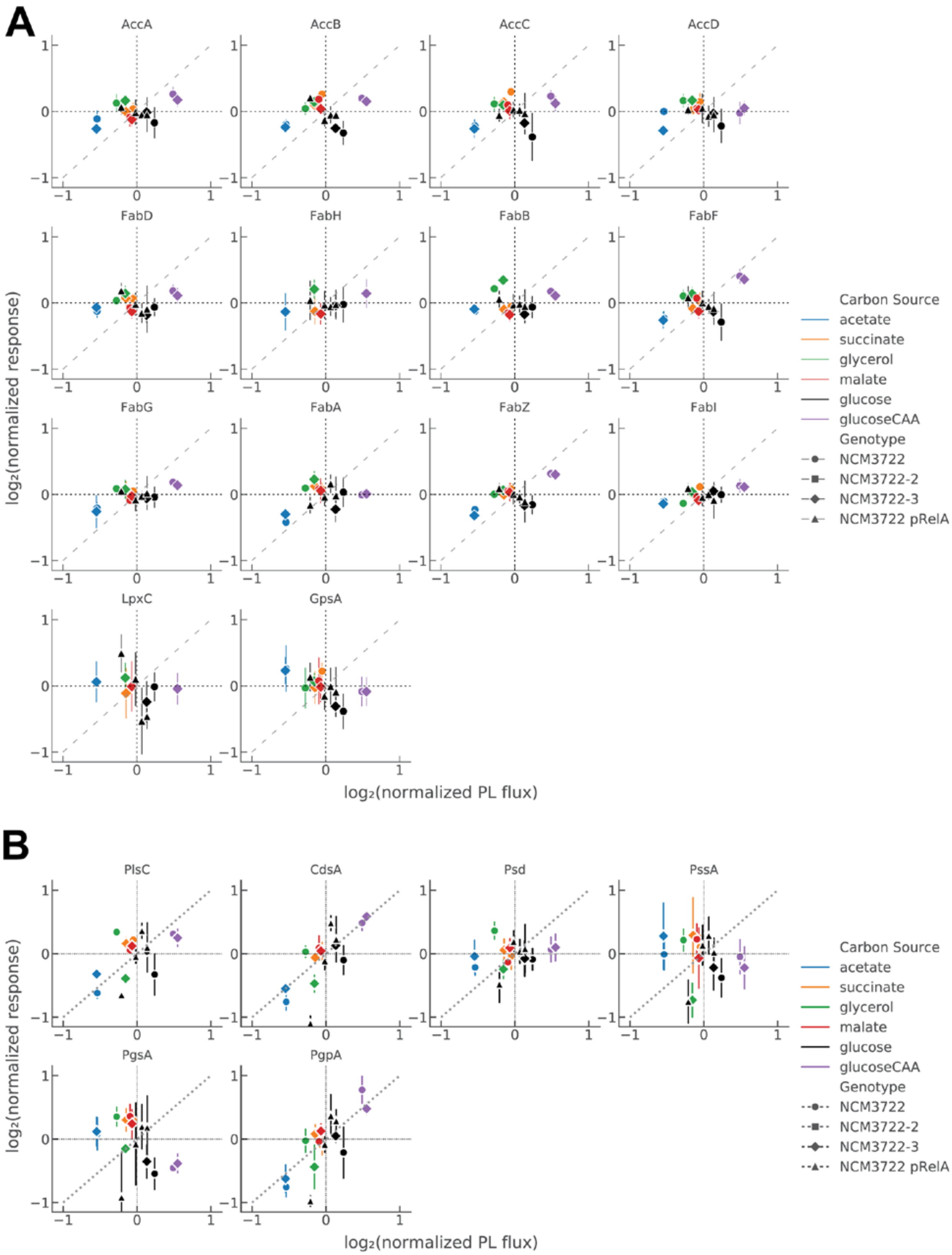
A
AccA
AccB
AccC
AccD
FabD
FabH
FabB
FabF
FabG
FabA
FabZ
FabI
LpxC
GpsA
log₂(normalized response)
log₂(normalized PL flux)
Carbon Source
acetate
succinate
glycerol
malate
glucose
glucoseCAA
Genotype
NCM3722
NCM3722-2
NCM3722-3
NCM3722 pRelA
B
PlsC
CdsA
Psd
PssA
PgsA
PgpA
log₂(normalized response)
log₂(normalized PL flux)
Carbon Source
acetate
succinate
glycerol
malate
glucose
glucoseCAA
Genotype
NCM3722
NCM3722-2
NCM3722-3
NCM3722 pRelA

**Supplemental Figure 2.** Steady-state concentrations of fatty acid and PL synthesis enzymes of *E. coli* NCM3722 in six media and of *E. coli* pRelA* in glucose cultures with titrated RelA* expression (triangles). Concentrations and PL fluxes are normalized to average across each individual replicate series. Error bars represent standard deviation of 3 sampling replicates. Two series of independent biological replicates were obtained for *E. coli* NCM3722.

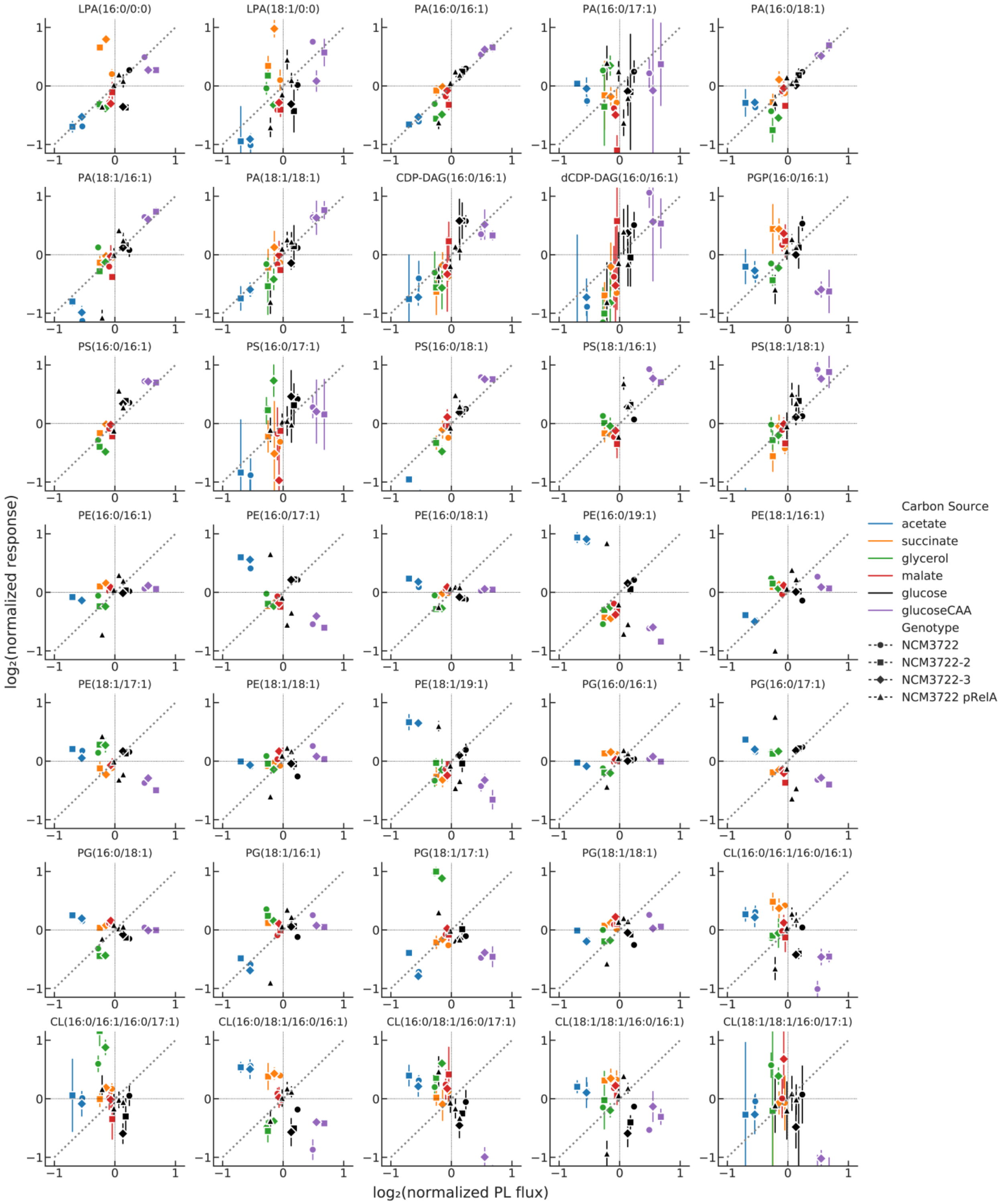


**Supplemental Figure 3.** Steady-state concentrations of PL synthesis intermediates of *E. coli* NCM3722 in six media and of *E. coli* pRelA* in glucose cultures with titrated RelA* expression (triangles). Concentrations and PL fluxes are normalized to average across each individual replicate series. Error bars represent standard deviation of 3 sampling replicates. Three series of independent biological replicates were obtained for *E. coli* NCM3722.

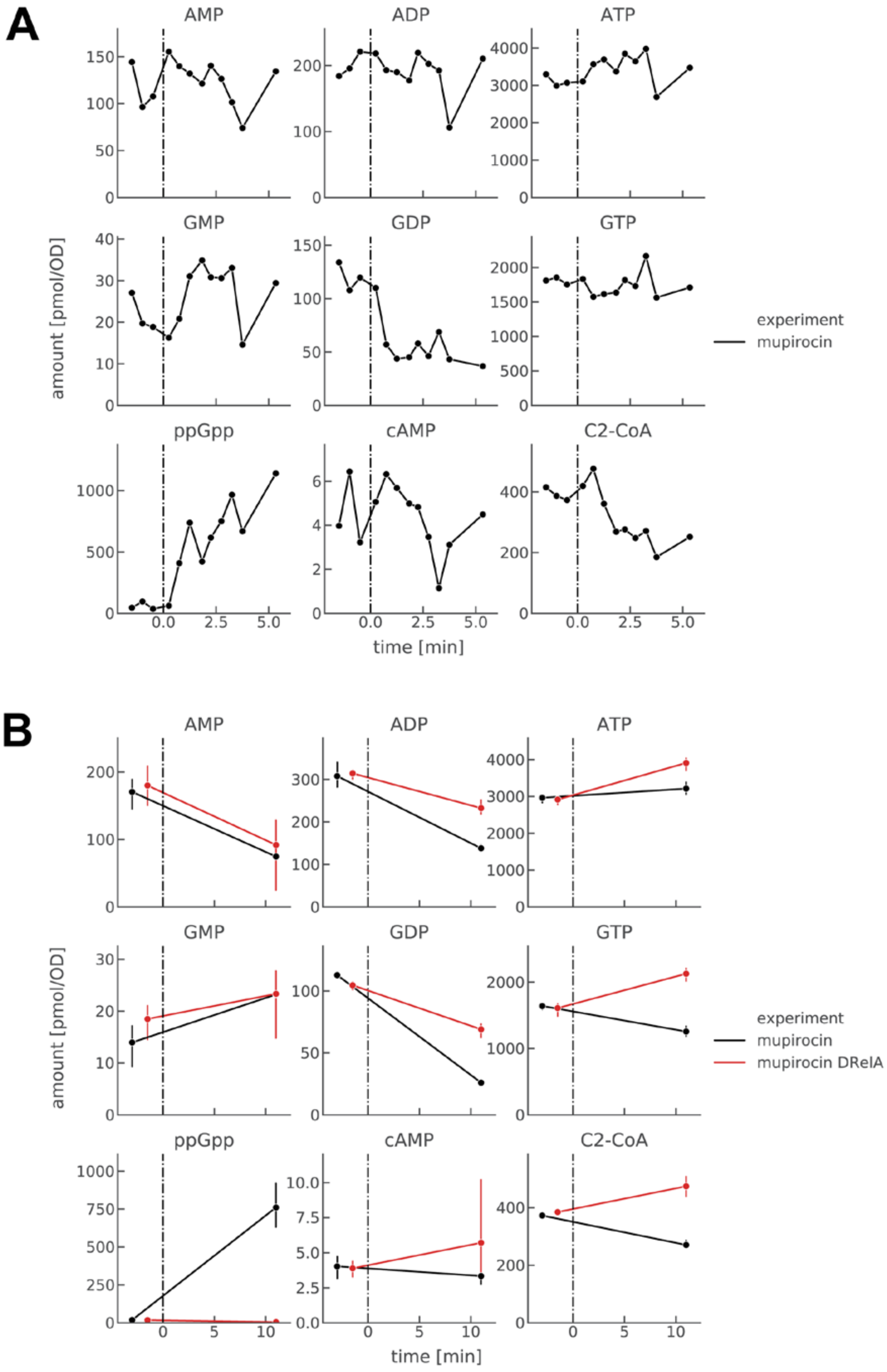
A
B
AMP
ADP
ATP
GMP
GDP
GTP
ppGpp
cAMP
C2-CoA
amount [pmol/OD]
time [min]
experiment
mupirocin
mupirocin DRelA

**Supplemental Figure 4. A.** Dynamics of nucleotide pools of wild-type *E. coli* after mupirocin addition (dashed line at t=0 minutes). Data points indicate individual measurements during a single time series. **B.** Response of nucleotide pools of wild-type and Δ*relA* strains to mupirocin, added at t=0. Error bars represent standard deviation of 3 sampling replicates from individual time series.

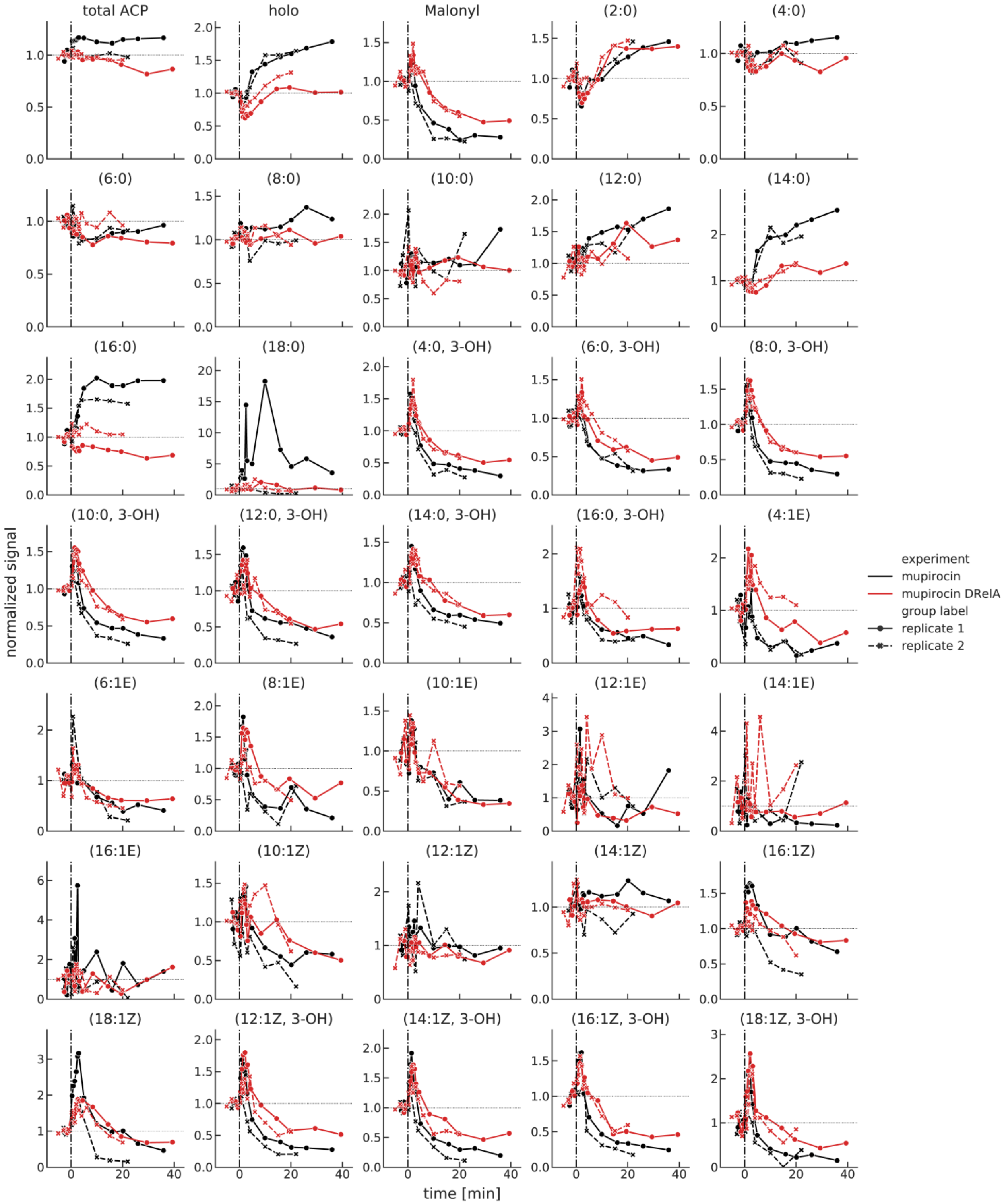


**Supplemental Figure 5.** Response of acyl-ACP species (counts per OD unit) to mupirocin at t = 0 in wild-type and Δ*relA E. coli.* Values are normalized such that the concentrations before t = 0

are averaged to 1. Data points indicate individual measurements from a single time series. Two independent biological replicates per strain are shown.

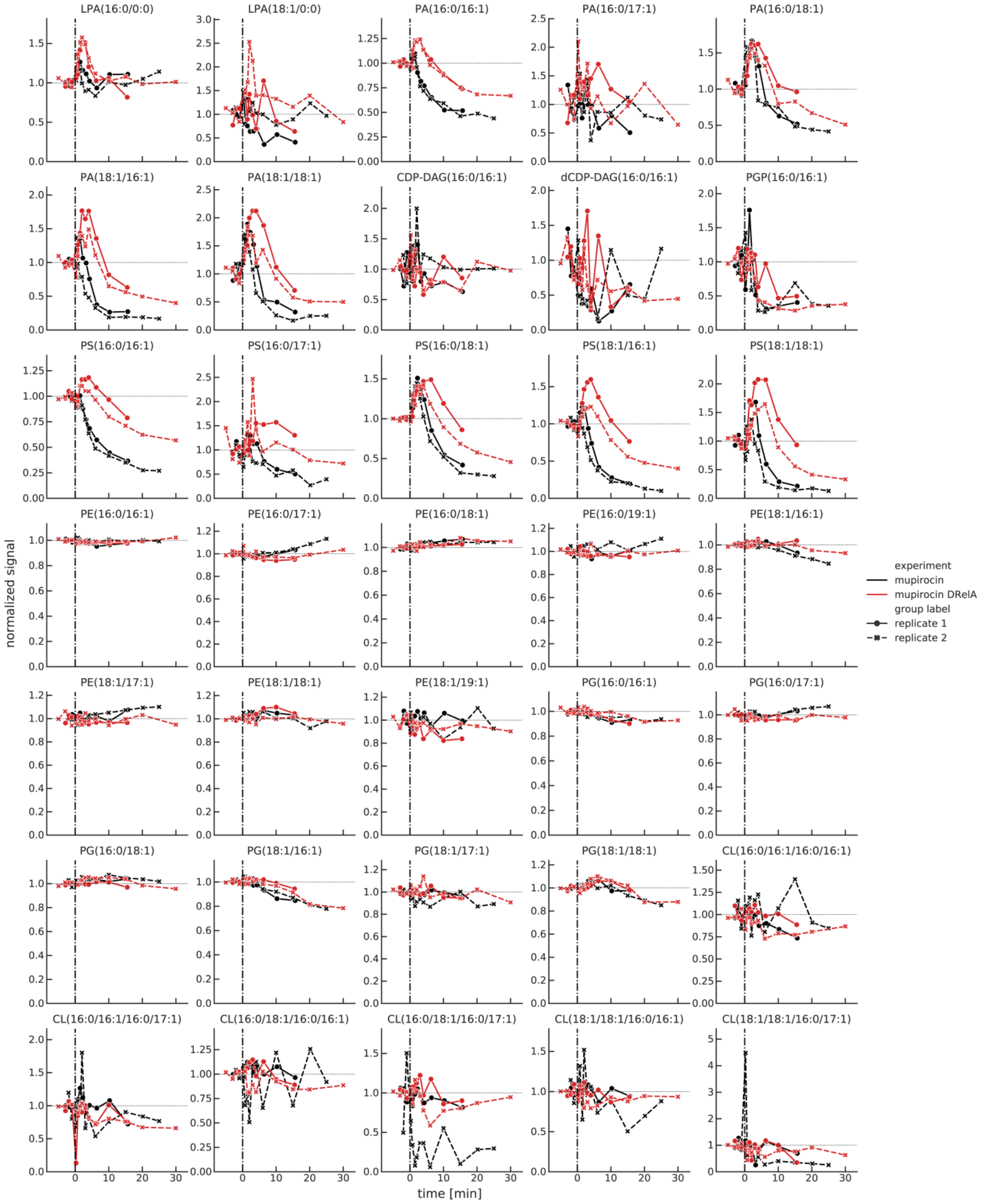


**Supplemental Figure 6.** Response of individual PL species (counts per total PE) to mupirocin at t = 0 in wild-type and Δ*relA E. coli.* Values are normalized such that the concentrations before t =

0 are averaged to 1. Data points indicate individual measurements from a single time series.

Two independent biological replicates per strain are shown.

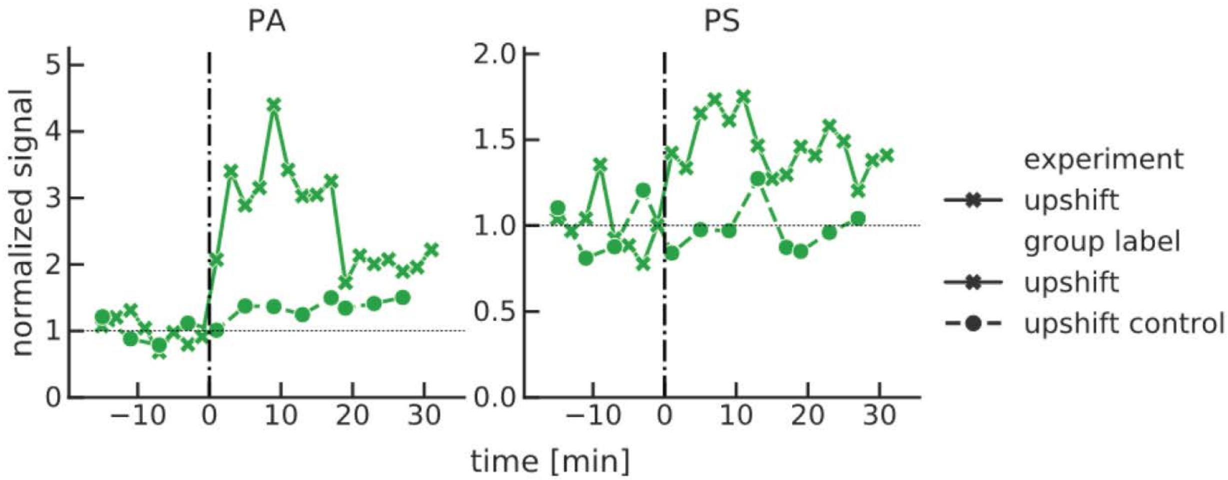


**Supplemental Figure 7.** Response of PL species (counts per total PE) to addition of glucose and amino acids at t = 0 to a glycerol culture of wild-type *E. coli.* Values are normalized such that the concentrations before t = 0 are averaged to 1. Data points indicate individual measurements from a single time series.

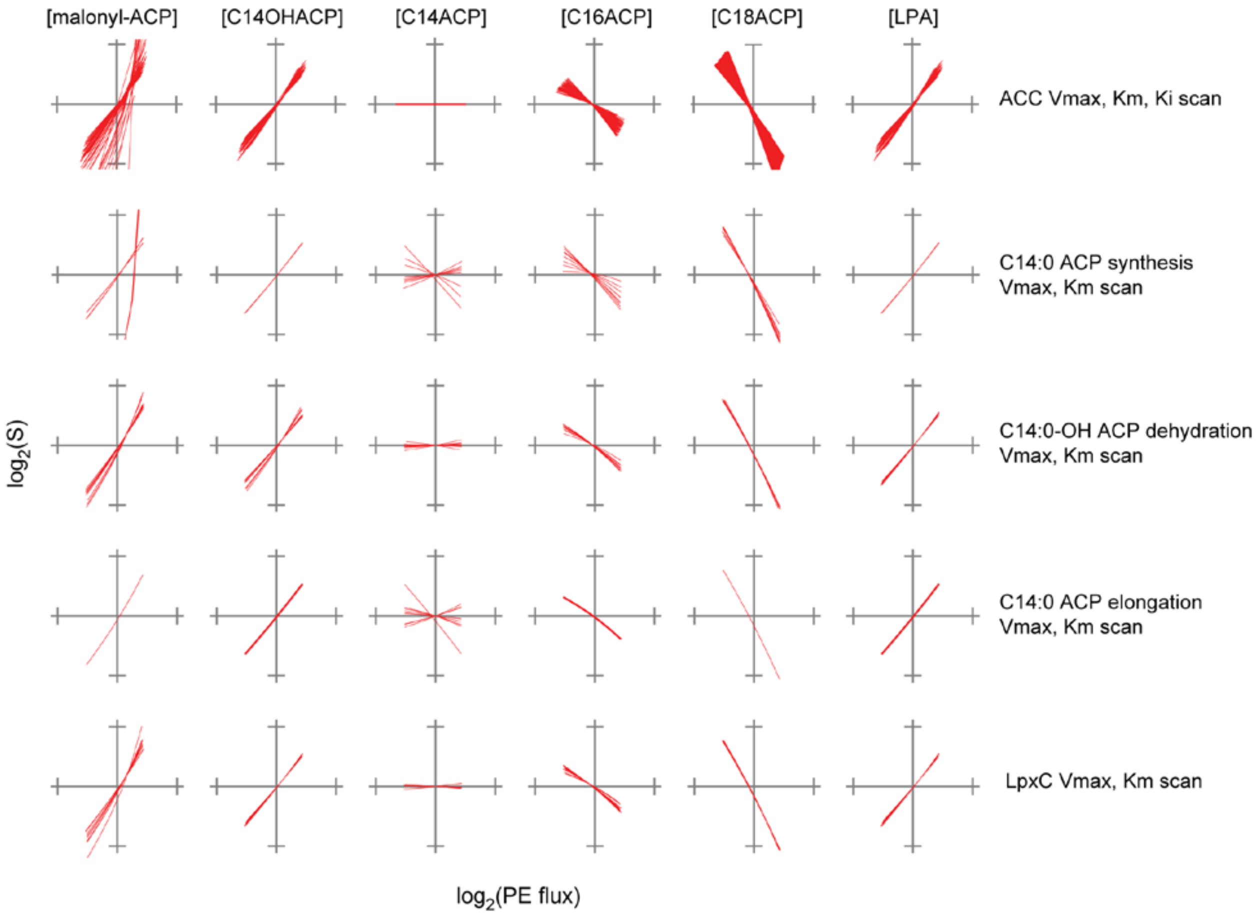


**Supplemental Figure 8.** Sensitivity analysis of the fatty acid and phospholipid synthesis model demonstrates that the steady-state metabolite concentration trends simulated from PlsB $V_{max}$ variations are robust against variations in model parameters. Parameters were simultaneously varied across a 4-fold range centered around the model parameter used. PE flux was varied by changing PlsB $V_{max}$ as described in the main text.

$$\frac{\mathrm{d}([\text{"malonyl-ACP"}] \cdot V_{\text{compartment}})}{\mathrm{d}t} = +V_{\text{compartment}} \cdot \left( \frac{V_{(\text{"acetyl-CoA carboxylase"})} \cdot [\text{"acetyl-CoA"}]}{[\text{"acetyl-CoA"}] + \mathrm{Km}_{(\text{"acetyl-CoA carboxylase"})} \cdot \left(1 + \frac{[\text{C16ACP}]}{\mathrm{Ki1}_{(\text{"acetyl-CoA carboxylase"})}} + \frac{[\text{C18ACP}]}{\mathrm{Ki2}_{(\text{"acetyl-CoA carboxylase"})}}\right)} \right)$$

$$-V_{\text{compartment}} \cdot \left( \frac{V_{(\text{"C14 synthesis"})} \cdot [\text{"malonyl-ACP"}]}{\mathrm{Km}_{(\text{"C14 synthesis"})} + [\text{"malonyl-ACP"}]} \right)$$

$$-V_{\text{compartment}} \cdot$$

$$\left( \frac{\mathrm{vmax}_{(\text{"C14 elongation"})} \cdot [\text{C14ACP}] \cdot [\text{"malonyl-ACP"}]}{\mathrm{Kma}_{(\text{"C14 elongation"})} \cdot \mathrm{Kmb}_{(\text{"C14 elongation"})} + [\text{C14ACP}] \cdot \mathrm{Kmb}_{(\text{"C14 elongation"})} + [\text{"malonyl-ACP"}] \cdot \mathrm{Kma}_{(\text{"C14 elongation"})} + [\text{C14ACP}] \cdot [\text{"malonyl-ACP"}]} \right)$$

$$-V_{\text{compartment}} \cdot$$

$$\left( \frac{\mathrm{vmax}_{(\text{"C16 elongation"})} \cdot [\text{C16ACP}] \cdot [\text{"malonyl-ACP"}]}{\mathrm{Kma}_{(\text{"C16 elongation"})} \cdot \mathrm{Kmb}_{(\text{"C16 elongation"})} + [\text{C16ACP}] \cdot \mathrm{Kmb}_{(\text{"C16 elongation"})} + [\text{"malonyl-ACP"}] \cdot \mathrm{Kma}_{(\text{"C16 elongation"})} + [\text{C16ACP}] \cdot [\text{"malonyl-ACP"}]} \right)$$

$$\frac{\mathrm{d}([\text{C16ACP}] \cdot V_{\text{compartment}})}{\mathrm{d}t} = +V_{\text{compartment}} \cdot \left( \frac{V_{(\text{"C16 dehydration"})} \cdot [\text{C16OHACP}]}{\mathrm{Km}_{(\text{"C16 dehydration"})} + [\text{C16OHACP}]} \right)$$

$$-V_{\text{compartment}} \cdot$$

$$\left( \frac{\mathrm{vmax}_{(\text{"C16 elongation"})} \cdot [\text{C16ACP}] \cdot [\text{"malonyl-ACP"}]}{\mathrm{Kma}_{(\text{"C16 elongation"})} \cdot \mathrm{Kmb}_{(\text{"C16 elongation"})} + [\text{C16ACP}] \cdot \mathrm{Kmb}_{(\text{"C16 elongation"})} + [\text{"malonyl-ACP"}] \cdot \mathrm{Kma}_{(\text{"C16 elongation"})} + [\text{C16ACP}] \cdot [\text{"malonyl-ACP"}]} \right)$$

$$-V_{\text{compartment}} \cdot \left( \frac{\mathrm{kcat}_{(\text{"C16 LPA synthesis"})} \cdot [\text{PlsB}] \cdot [\text{C16ACP}]}{\mathrm{Km}_{(\text{"C16 LPA synthesis"})} + [\text{C16ACP}]} \right)$$

$$\frac{\mathrm{d}([\text{LPA}] \cdot V_{\text{compartment}})}{\mathrm{d}t} = +V_{\text{compartment}} \cdot \left( \frac{\mathrm{kcat}_{(\text{"C18 LPA synthesis"})} \cdot [\text{PlsB}] \cdot [\text{C18ACP}]}{\mathrm{Km}_{(\text{"C18 LPA synthesis"})} + [\text{C18ACP}]} \right)$$

$$+V_{\text{compartment}} \cdot \left( \frac{\mathrm{kcat}_{(\text{"C16 LPA synthesis"})} \cdot [\text{PlsB}] \cdot [\text{C16ACP}]}{\mathrm{Km}_{(\text{"C16 LPA synthesis"})} + [\text{C16ACP}]} \right)$$

$$-V_{\text{compartment}} \cdot \left( \frac{\mathrm{vmax}_{(\text{"PA synthesis"})} \cdot [\text{LPA}] \cdot [\text{C161ACP}]}{\mathrm{Kma}_{(\text{"PA synthesis"})} \cdot \mathrm{Kmb}_{(\text{"PA synthesis"})} + [\text{LPA}] \cdot \mathrm{Kmb}_{(\text{"PA synthesis"})} + [\text{C161ACP}] \cdot \mathrm{Kma}_{(\text{"PA synthesis"})} + [\text{LPA}] \cdot [\text{C161ACP}]} \right)$$

$$\frac{\mathrm{d}([\text{PA}] \cdot V_{\text{compartment}})}{\mathrm{d}t} = +V_{\text{compartment}} \cdot \left( \frac{\mathrm{vmax}_{(\text{"PA synthesis"})} \cdot [\text{LPA}] \cdot [\text{C161ACP}]}{\mathrm{Kma}_{(\text{"PA synthesis"})} \cdot \mathrm{Kmb}_{(\text{"PA synthesis"})} + [\text{LPA}] \cdot \mathrm{Kmb}_{(\text{"PA synthesis"})} + [\text{C161ACP}] \cdot \mathrm{Kma}_{(\text{"PA synthesis"})} + [\text{LPA}] \cdot [\text{C161ACP}]} \right)$$

$$-V_{\text{compartment}} \cdot \left( \frac{V_{(\text{"CDPDAG synthesis"})} \cdot [\text{PA}]}{\mathrm{Km}_{(\text{"CDPDAG synthesis"})} + [\text{PA}]} \right)$$

$$\frac{\mathrm{d}([\text{CDPDAG}] \cdot V_{\text{compartment}})}{\mathrm{d}t} = +V_{\text{compartment}} \cdot \left( \frac{V_{(\text{"CDPDAG synthesis"})} \cdot [\text{PA}]}{\mathrm{Km}_{(\text{"CDPDAG synthesis"})} + [\text{PA}]} \right)$$

$$-V_{\text{compartment}} \cdot \left( \frac{V_{(\text{"PS synthesis"})} \cdot [\text{CDPDAG}]}{\mathrm{Km}_{(\text{"PS synthesis"})} + [\text{CDPDAG}]} \right)$$

$$\frac{\mathrm{d}([\mathrm{PS}]\cdot V_{\mathrm{compartment}})}{\mathrm{d}t} = +V_{\mathrm{compartment}}\cdot\left(\frac{V_{(\text{"PS synthesis"})}\cdot[\mathrm{CDPDAG}]}{\mathrm{Km}_{(\text{"PS synthesis"})}+[\mathrm{CDPDAG}]}\right)$$

$$-V_{\mathrm{compartment}}\cdot\left(\frac{V_{(\text{"PE synthesis"})}\cdot[\mathrm{PS}]}{\mathrm{Km}_{(\text{"PE synthesis"})}+[\mathrm{PS}]}\right)$$

$$\frac{\mathrm{d}([\mathrm{C14BKACP}]\cdot V_{\mathrm{compartment}})}{\mathrm{d}t} = +V_{\mathrm{compartment}}\cdot\left(\frac{V_{(\text{"C14 synthesis"})}\cdot[\text{"malonyl}-\text{ACP"}]}{\mathrm{Km}_{(\text{"C14 synthesis"})}+[\text{"malonyl}-\text{ACP"}]}\right)$$

$$-V_{\mathrm{compartment}}\cdot\left(\frac{V_{(\text{"C14 reduction"})}\cdot[\mathrm{C14BKACP}]}{\mathrm{Km}_{(\text{"C14 reduction"})}+[\mathrm{C14BKACP}]}\right)$$

$$\frac{\mathrm{d}([\mathrm{C14OHACP}]\cdot V_{\mathrm{compartment}})}{\mathrm{d}t} = +V_{\mathrm{compartment}}\cdot\left(\frac{V_{(\text{"C14 reduction"})}\cdot[\mathrm{C14BKACP}]}{\mathrm{Km}_{(\text{"C14 reduction"})}+[\mathrm{C14BKACP}]}\right)$$

$$-V_{\mathrm{compartment}}\cdot\left(\frac{V_{(\text{"LPS initiation"})}\cdot[\mathrm{C14OHACP}]}{\mathrm{Km}_{(\text{"LPS initiation"})}+[\mathrm{C14OHACP}]}\right)$$

$$-V_{\mathrm{compartment}}\cdot\left(\frac{V_{(\text{"C14 dehydration"})}\cdot[\mathrm{C14OHACP}]}{\mathrm{Km}_{(\text{"C14 dehydration"})}+[\mathrm{C14OHACP}]}\right)$$

$$\frac{\mathrm{d}([\mathrm{C14ACP}]\cdot V_{\mathrm{compartment}})}{\mathrm{d}t} = -V_{\mathrm{compartment}}\cdot$$

$$\left(\frac{\mathrm{vmax}_{(\text{"C14 elongation"})}\cdot[\mathrm{C14ACP}]\cdot[\text{"malonyl}-\text{ACP"}]}{\mathrm{Kma}_{(\text{"C14 elongation"})}\cdot\mathrm{Kmb}_{(\text{"C14 elongation"})}+[\mathrm{C14ACP}]\cdot\mathrm{Kmb}_{(\text{"C14 elongation"})}+[\text{"malonyl}-\text{ACP"}]\cdot\mathrm{Kma}_{(\text{"C14 elongation"})}+[\mathrm{C14ACP}]\cdot[\text{"malonyl}-\text{ACP"}]}\right)$$

$$+V_{\mathrm{compartment}}\cdot\left(\frac{V_{(\text{"C14 dehydration"})}\cdot[\mathrm{C14OHACP}]}{\mathrm{Km}_{(\text{"C14 dehydration"})}+[\mathrm{C14OHACP}]}\right)$$

$$\frac{\mathrm{d}([\mathrm{C16BKACP}]\cdot V_{\mathrm{compartment}})}{\mathrm{d}t} = +V_{\mathrm{compartment}}\cdot$$

$$\left(\frac{\mathrm{vmax}_{(\text{"C14 elongation"})}\cdot[\mathrm{C14ACP}]\cdot[\text{"malonyl}-\text{ACP"}]}{\mathrm{Kma}_{(\text{"C14 elongation"})}\cdot\mathrm{Kmb}_{(\text{"C14 elongation"})}+[\mathrm{C14ACP}]\cdot\mathrm{Kmb}_{(\text{"C14 elongation"})}+[\text{"malonyl}-\text{ACP"}]\cdot\mathrm{Kma}_{(\text{"C14 elongation"})}+[\mathrm{C14ACP}]\cdot[\text{"malonyl}-\text{ACP"}]}\right)$$

$$-V_{\mathrm{compartment}}\cdot\left(\frac{V_{(\text{"C16 reduction"})}\cdot[\mathrm{C16BKACP}]}{\mathrm{Km}_{(\text{"C16 reduction"})}+[\mathrm{C16BKACP}]}\right)$$

$$\frac{\mathrm{d}([\mathrm{C16OHACP}]\cdot V_{\mathrm{compartment}})}{\mathrm{d}t} = +V_{\mathrm{compartment}}\cdot\left(\frac{V_{(\text{"C16 reduction"})}\cdot[\mathrm{C16BKACP}]}{\mathrm{Km}_{(\text{"C16 reduction"})}+[\mathrm{C16BKACP}]}\right)$$

$$-V_{\mathrm{compartment}}\cdot\left(\frac{V_{(\text{"C16 dehydration"})}\cdot[\mathrm{C16OHACP}]}{\mathrm{Km}_{(\text{"C16 dehydration"})}+[\mathrm{C16OHACP}]}\right)$$

$$\frac{\mathrm{d}([\mathrm{C18BKACP}]\cdot V_{\mathrm{compartment}})}{\mathrm{d}t} = +V_{\mathrm{compartment}}\cdot$$

$$\left(\frac{\mathrm{vmax}_{(\text{"C16 elongation"})}\cdot[\mathrm{C16ACP}]\cdot[\text{"malonyl}-\text{ACP"}]}{\mathrm{Kma}_{(\text{"C16 elongation"})}\cdot\mathrm{Kmb}_{(\text{"C16 elongation"})}+[\mathrm{C16ACP}]\cdot\mathrm{Kmb}_{(\text{"C16 elongation"})}+[\text{"malonyl}-\text{ACP"}]\cdot\mathrm{Kma}_{(\text{"C16 elongation"})}+[\mathrm{C16ACP}]\cdot[\text{"malonyl}-\text{ACP"}]}\right)$$

$$-V_{\mathrm{compartment}}\cdot\left(\frac{V_{(\text{"C18 reduction"})}\cdot[\mathrm{C18BKACP}]}{\mathrm{Km}_{(\text{"C18 reduction"})}+[\mathrm{C18BKACP}]}\right)$$

$$\frac{\mathrm{d}([\mathrm{C18OHACP}] \cdot V_{\mathrm{compartment}})}{\mathrm{d}t} = +V_{\mathrm{compartment}} \cdot \left(\frac{V_{(\text{"C18 reduction"})} \cdot [\mathrm{C18BKACP}]}{\mathrm{Km}_{(\text{"C18 reduction"})} + [\mathrm{C18BKACP}]}\right) - V_{\mathrm{compartment}} \cdot \left(\frac{V_{(\text{"C18 dehydration"})} \cdot [\mathrm{C18OHACP}]}{\mathrm{Km}_{(\text{"C18 dehydration"})} + [\mathrm{C18OHACP}]}\right)$$

$$\frac{\mathrm{d}([\mathrm{C18ACP}] \cdot V_{\mathrm{compartment}})}{\mathrm{d}t} = +V_{\mathrm{compartment}} \cdot \left(\frac{V_{(\text{"C18 dehydration"})} \cdot [\mathrm{C18OHACP}]}{\mathrm{Km}_{(\text{"C18 dehydration"})} + [\mathrm{C18OHACP}]}\right) - V_{\mathrm{compartment}} \cdot \left(\frac{\mathrm{kcat}_{(\text{"C18 LPA synthesis"})} \cdot [\mathrm{PlsB}] \cdot [\mathrm{C18ACP}]}{\mathrm{Km}_{(\text{"C18 LPA synthesis"})} + [\mathrm{C18ACP}]}\right)$$

**Supplemental Table 1.** Differential equations used to define steady-state fluxes in the model. Note that ppGpp inhibition of PlsB was not considered in steady-state calculations (i.e. ppGpp was set to 0 μM).

**Supplemental Text 1.**

**Steady-state model description**

The simplified pathway model was set up using COPASI as a series of irreversible Michaelis-Menten equations provided in **Supplemental Table 1**. Most reactions in the fatty acid pathway were excluded in order to determine whether the experimentally-observed trends in intermediate concentrations could be captured in a simplified model. Acetyl-CoA and C16:1-ACP concentrations were fixed at their initial values. For simplicity, $V_{max}$ values and $K_M$ parameters of each reaction were set to similar values (around 10 and 30 µM, full parameter set given below). For steady-state calculations, we do not claim that the model parameters reflect the exact *in vivo* values. However the values are useful in capturing the basic steady-state behaviours of the system. The simulated steady-state fluxes and metabolite concentrations depicted in **Figure 2** were obtained using the “Parameter Scan” function in COPASI. The differential equations that determine the fluxes through each intermediate pool are defined below.

The strong flux control exerted by PlsB is a consequence of end-product inhibition of ACC by C16:0-ACP and C18:0-ACP, as predicted by metabolic control analysis. Although acyl-ACP species are predicted from experiments to exhibit mixed inhibition of *E. coli* ACC with respect to acetyl-CoA, we use competitive inhibition to minimize the number of parameters in the model. Changes in concentrations of other substrates (e.g. ATP, bicarbonate) or allosteric regulators of ACC (e.g. GlnB) would be expected to exert a similar influence on ACC activity and fatty acid synthesis as variations in ACC $V_{max}$.

| Values used in mathematical model of fatty acid and PL synthesis pathways. | | | |
|---|---|---|---|
| **Initial Species Values** | Species | | Initial Concentration |
| | PlsB | | 1 µmol/l (varied) |
| | acetyl-CoA | | 300 µmol/l (fixed) |
| | malonyl-ACP | | 0 µmol/l |
| | C16ACP | | 0 µmol/l |
| | C18ACP | | 0 µmol/l |
| | LPA | | 0 µmol/l |
| | ppGpp | | 0 µmol/l |
| | PA | | 0 µmol/l |
| | CDPDAG | | 0 µmol/l |
| | PS | | 0 µmol/l |
| | C161ACP | | 30 µmol/l (fixed) |
| | C14BKACP | | 0 µmol/l |
| | C14OHACP | | 0 µmol/l |
| | C14ACP | | 0 µmol/l |
| | C16BKACP | | 0 µmol/l |
| | C16OHACP | | 0 µmol/l |
| | C18BKACP | | 0 µmol/l |
| | C18OHACP | | 0 µmol/l |
| **Kinetic Parameters** | Reaction | Parameter | Value |
| | acetyl-CoA carboxylase | | |
| | | Km | 300 µmol |
| | | V | 50 µmol/s |
| | | Ki1 | 1 µmol |
| | | Ki2 | 1 µmol |
| | C14 synthesis | | |
| | | Km | 30 µmol |
| | | V | 10 µmol/s |
| | C16 LPA synthesis | | |
| | | Km | 30 µmol |
| | | kcat | 10 µmol/s |
| | PA synthesis | | |
| | | vmax | 20 µmol/s |
| | | Kma | 30 µmol |
| | | Kmb | 30 µmol |
| | CDPDAG synthesis | | |
| | | Km | 30 µmol |
| | | V | 10 µmol/s |
| | PS synthesis | | |
| | | Km | 30 µmol |
| | | V | 10 µmol/s |
| | PE synthesis | | |
| | | Km | 30 µmol |
| | | V | 10 µmol/s |
| | C14 reduction | | |
| | | Km | 30 µmol |
| | | V | 10 µmol/s |
| | LPS initiation | | |
| | | Km | 30 µmol |
| | | V | 10 µmol/s |
| | C14 dehydration | | |

| | | |
|---|---|---|
| | Km | 30 µmol |
| | V | 10 µmol/s |
| C14 elongation | | |
| | vmax | 20 µmol/s |
| | Kma | 30 µmol |
| | Kmb | 30 µmol |
| C16 reduction | | |
| | Km | 30 µmol |
| | V | 10 µmol/s |
| C16 dehydration | | |
| | Km | 30 µmol |
| | V | 10 µmol/s |
| C16 elongation | | |
| | vmax | 20 µmol/s |
| | Kma | 30 µmol |
| | Kmb | 30 µmol |
| C18 reduction | | |
| | Km | 30 µmol |
| | V | 10 µmol/s |
| C18 dehydration | | |
| | Km | 30 µmol |
| | V | 10 µmol/s |
| C18 LPA synthesis | | |
| | Km | 30 µmol |
| | kcat | 10 µmol/s |
